# Aqueous two-phase bioinks for discrete packing and compartmentalisation of 3D bioprinted cells

**DOI:** 10.1101/2025.06.27.661968

**Authors:** Martina Marcotulli, Arianna Iacomino, Federico Serpe, Lucia Iafrate, Marco Bastioli, Giorgia Montalbano, Biagio Palmisano, Silvia Franco, Roberta Angelini, Alessandro Corsi, Mara Riminucci, Giancarlo Ruocco, Chiara Scognamiglio, Andrea Barbetta, Gianluca Cidonio

## Abstract

The unparalleled ability of aqueous two-phase systems (ATPS) to reproduce microscale cellular and biomaterial compartmentalisation to selectively modulate cell behaviour and functionality is ideal for tissue engineering and regenerative medicine (TERM) purposes. Herein, we introduce new ATPS biomaterial inks for 3D bioprinting of water-in-water (W/W) emulsions, enabling precise cellular crowding for tissue regeneration *in vitro* and *ex vivo*. Gelatin methacryloyl (GelMA) was hierarchically dispersed in an alginic acid phase depending on sodium chloride (NaCl) concentration (0-36 g/L). Emulsion droplet size (12.8±2.6 µm to 52.4±11.4 µm) influenced degradation and spatial cell localisation (A549, C2C12, MG63). A microfluidic-assisted 3D bioprinting approach allowed fine-tuning of fibre structure adjusting ATPS deposition by modulating flow rates and printing speed. Rheological properties supported the findings of the two-phase partitioning, aiding the selection of the ATPS ink formulation for functional cell-laden construct fabrication. Encapsulation of C2C12 cells revealed enhanced cytoskeletal remodelling at higher salt concentrations. Increased GelMA phase promoted human bone marrow stromal cells (HBMSCs) crowding, mineral deposition and skeletal differentiation. *In ovo* studies demonstrated degradation control and vascular infiltration via salt modulation. Altogether, ATPS bioinks offer a versatile platform for the assembling of complex, hierarchical tissues with microscale precision, expanding biofabrication strategies for TERM applications.

## 1. Introduction

Aqueous two-phase systems (ATPS) are liquid-liquid partitioning systems mainly used for separating, purifying, and enriching biomolecules such as proteins, viruses, and enzymes.[1] These systems are typically engineered from two water-soluble components, such as polymers, salts, alcohols or ionic liquids, which are mixed in water at sufficient concentrations, leading to the formation of two immiscible aqueous phases.[2] Each phase predominantly contains one of the materials, creating an ideal environment for various bioseparation processes.[3]

Contrarily to oil-in-water (O/W) systems, ATPS can be engineered to sustain cellular encapsulation, thus facilitating biological functionality and stability for bioengineering purposes. Common phase-forming agents include polymers like polyethylene glycol (PEG) and dextran,[4,5] or a combination of PEG and salts like phosphate,[6] sulfate,[7] or citrate.[8] The phase separation is driven by thermodynamic forces, where the enthalpic interactions between the polymers and water molecules lead to the formation of two distinct aqueous phases. The process balances the enthalpy and entropy of the system, resulting in immiscible liquid compartments.[9] Nowadays, the use of ATPS could represent an emerging strategy in tissue engineering and regenerative medicine (TERM), which typically aims to create biological substitutes to restore, sustain or enhance the functionality of damaged tissues. Indeed, ATPS could be modulated in concentrations to enable the spatial organisation of cells and biomolecules via phase compartmentalisation. This level of control is essential to replicate natural native tissue environments and new models for tissue regeneration.[10] Moreover, their aqueous composition allows ATPS to demonstrate biocompatible properties for biological applications. In fact, ATPS systems have been found able to support cell viability, proliferation and differentiation of living cells.[11] Among recent applications, ATPS have been used to develop new bioinks in 3D bioprinting.[12] The water-based inks provide precise deposition of cells and biomaterials, enabling the creation of complex tissue structures and maintaining high cell viability. In addition, the ability of ATPS to compartmentalise cells and biomolecules components into distinct microenvironments allows unique opportunities for mimicking the heterogeneous architecture of tissues.[13] In a pioneering work by Ying and co-workers[14] demonstrated a new ATPS ink from poly-ethylene oxide (PEO) and gelatin methacryloyl (GelMA)., with the internal PEO phase gradually diffusing out of the laden fibres once in culture, forming a porous structure that facilitated nutrient-waste exchange, driving cell migration and proliferation. Further findings focused on the tuning of pore size,[15] enhancing ATPS[16] stability, and on the use of different polymeric components to tune scaffold material properties and function.[17–20] However, an unsolved challenge currently limiting the use of ATPS in extrusion-based 3D bioprinting is the weak viscoelastic characteristics, which prevent direct writing. A number of strategies have been implemented to overcome this limitation such as adjusting the temperature close to the gelling temperature[14] (in the case of GelMA-based ATPS), patterning ATPS in a suspended medium,[18,21] or adding rheological modifiers to improve ATPS stability.[19] While resulting effective, the aforementioned strategies implemented for extrusion-based approaches resulted valid exclusively for the specific polymers present in the ATPS continuous phase. Digital light processing (DLP) platforms have been employed to overcome the viscoelastic requirements,[18,21] while suffering from recurrent drawbacks such as the waste of uncross-linked ATPS ink and photo-toxicity. Recently, microfluidic-assisted 3D bioprinting strategies have been proposed to drive the controlled extrusion of biphasic inks and the deposition of cellular bioinks for multi-tissue fabrication.[22,23] Thus, the synergistic use of ATPS ink with microfluidic-assisted 3D bioprinting strategy holds enormous potential to advance tissue-specific regeneration by enabling the precise spatial arrangement of different cell types, which is critical for functional tissue restoration.[12,24] Due to their biocompatibility and oil-free nature, ATPS include cells within stable microdroplets formed by the immiscible aqueous phases, preserving their functionality and integrity during encapsulation.[25] However, replicating biocompatible ATPS systems is often challenging and associated with a number of complications. A recent solution to guide cell compartmentalisation has been offered by a caseinate solution in-alginate to form a stable ATPS localising human white adipose progenitors[26] (WAPs) within the pores resulting from the evacuation of the dispersed phase. Although this approach was found to guide the adipogenic differentiation of WAPs, the simple processing via casting approach, greatly limited the possibility to generate three-dimensional functional constructs.

In this research work, we present a unique approach to create and 3D bioprint a new class of ATPS biomaterial inks, specifically designed for TERM applications. The system relies on the controlled mixing of two biomaterials (GelMA and alginic acid) with ATPS facilitated by salt solution of sodium chloride (NaCl) at varying concentrations (0-36 g/L) (**Figure 1**). We have here demonstrated the compartmentalisation of GelMA within continuous alginate phase for water-in-water (W/W) emulsion formation. The ability to tune the size of the dispersed phase was influenced by the regulation of NaCl concentrations, providing flexibility for controlling material ink degradation and cellular phenotype following 3D bioprinting. Building on a novel microfluidic-assisted 3D bioprinting platform, we have demonstrated the deposition of low viscoelastic ATPS inks in guiding the proliferation and function of a variety of cell types, including lung cancer, myoblasts, osteosarcoma and human bone marrow stromal cells (HBMSCs). We have demonstrated the possibility to modulate cellular crowding, by enhancing the GelMA dispersed phase, thus guiding HBMSCs towards osteogenic differentiation. Overall, the reported ATPS ink offers a promising templating platform, with compartmentalisation ability for cellular crowding, towards the engineering of complex tissue architectures with microscale precision, increasing the potential for engineering tissue substitutes with high biological functionality.

**Figure 1.**
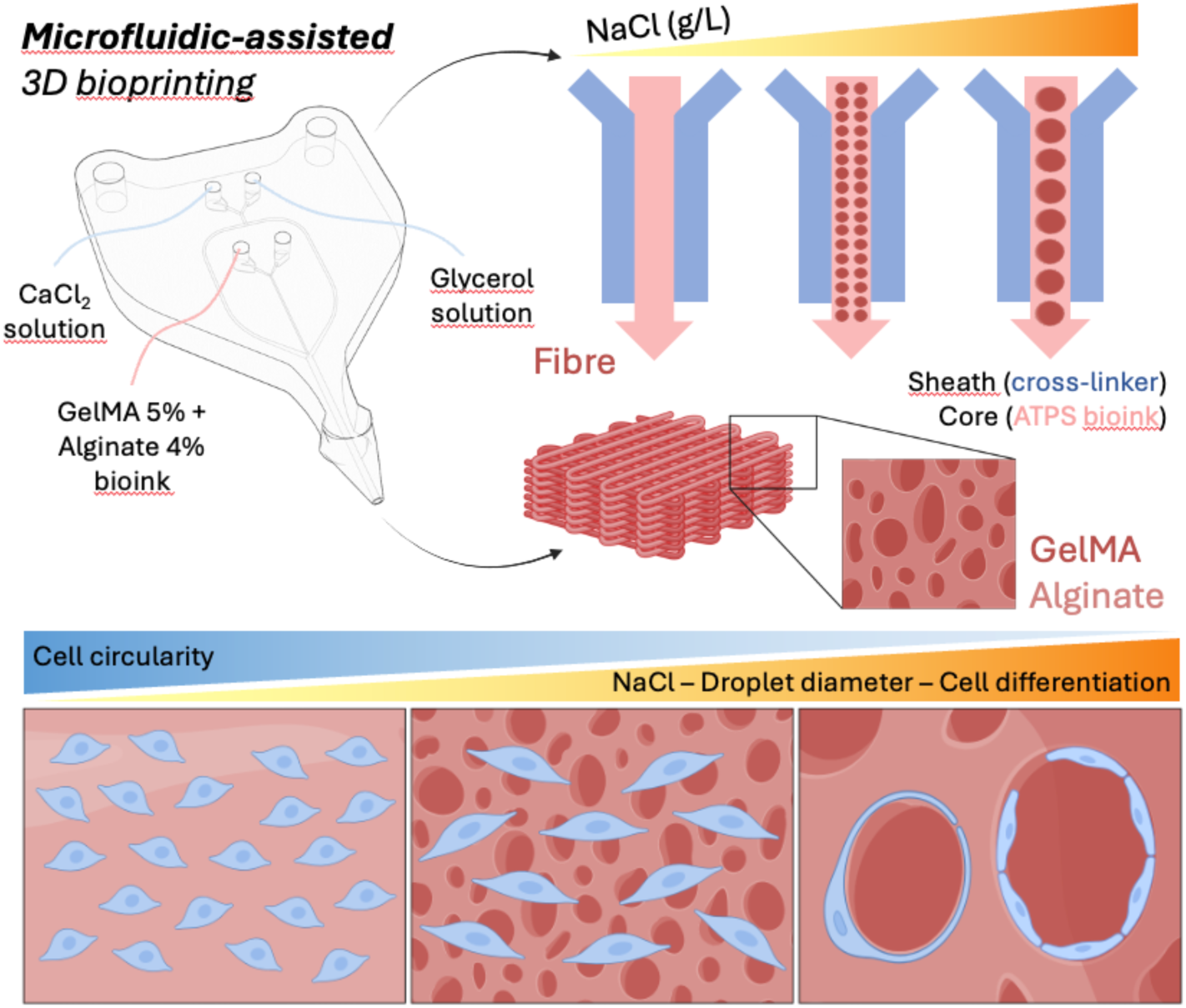
Rational of ATPS inks bioprinting. (left) The microfluidic printhead is integrated into a 3D bioprinting system to print the ATPS fibres. In the upper inlets, (i) a glycerol solution and (ii) CaCl_2_ cross-link solution are used. In the core inlet, ATPS inks based on GelMA 5% and Alginate 4% are flown. (right) The size of the ATPS droplets varies with increasing NaCl and GelMA concentration. (bottom) Within the ATPS hydrogel, cell organisation varies with droplet size, which in turn depends on NaCl concentration. From left to right, cells become increasingly elongated and organised within the GelMA droplets.

## 2. Results and discussion

### 2.1 Physical characterisation of ATPs inks demonstrated compartmentalised internal architecture

A library of ATPS formulation was prepared to observe the effect of salting out and the macroscopic behaviour of the emulsions. A marked increase in the dispersed phase volume was observed as evidenced by the progressive increase in the lower phase (GelMA) of the ATPS system. In particular, it was noted that in the absence of NaCl, no ATPS formation was observed, with a single-phase system and a completely transparent solution being observed within the vial (**Figure 2a**).

**Figure 2.**
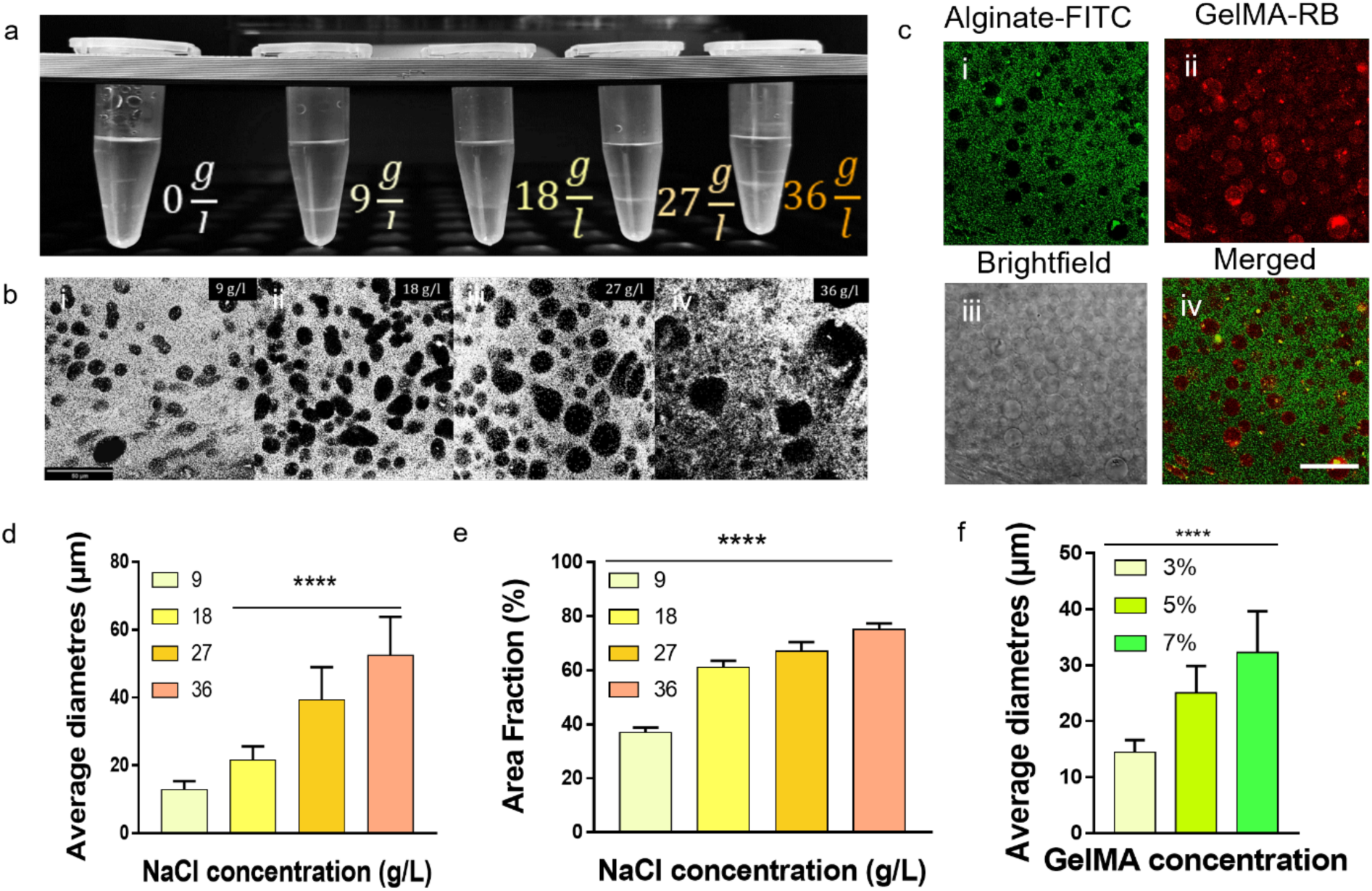
Physical and morphological analysis of ATPSs. a) Images of the different degrees of partitioning of the alginate- and GelMA-rich phases as the salt concentration increases. b) Brightfield confocal images of the ATPS as the NaCl content increases. In white, the external alginate phase; in black, the internal GelMA droplets. c) Confocal fluorescence images of the ATPS emulsion. c-i) External alginate phase labelled with FITC, c-ii) internal GelMA phase labelled with rhodamine B, c-iii) brightfield image of the ATPS, c-iv) merged fluorescence image. d) Average of droplet diameters measured at different salt concentrations. e) Volume fraction of dispersed phase calculated for ATPS at different NaCl concentrations. f) Average droplet diameters measured at different GelMA contents. Scale bars: (b) 50 μm, (c) 30 μm. Statistical significances were assessed by one-way ANOVA. Mean ± S.D. n=50, ****p<0.0001.

In contrast, increasing the salt percentage led to a progressively biphasic and opaque system. By confocal microscopy, the internal structure of the ATPS was validated, observing an increase in the volume fraction of the dispersed phase as the salt content increased (**Figure 2b**). The distribution of the two phases of alginate and GelMA was clarified by labelling alginate with FITC and GelMA with Rhodamine B. Following the formation of the ATPS, the structure was observed by confocal microscopy (**Figure 2c**) and the internal distribution of GelMA-RB and the external distribution of Alginate-FITC were confirmed. The size of the GelMA-rich phase droplets (**Figure 2d**) and the area fraction of the dispersed phase (**Figure 2e**) were found to be directly related to the salt content. Furthermore, a similar trend in significant proportionality was observed between the average size of the droplets and the GelMA content (**Figure 2f**).

These observations are consistent with previously reported behaviours in aqueous biphasic systems, where the increase in salt concentration increases the salting-out effect, promoting phase separation and the formation of larger droplets. With higher salt concentration, increased interfacial tension between phases was observed, which facilitates the coalescence of smaller droplets into larger droplets. In addition, the higher ionic strength was shown to reduce the electrostatic repulsion between the droplets, facilitating their fusion and contributing to the observed increase in dispersed phase.[27,28]

### 2.2 ATPs inks can be tuned by altering the salt and dispersed phase concentration

The tunability of ATPs inks through the manipulation of salt and GelMA concentrations was investigated, focusing on how these variables influence the size of the internal phase droplets. By examining different ATPS formulations, we provided detailed insights into the correlation between salt concentration, GelMA percentage and droplet size distribution. Microscopic images of the emulsions were obtained by flowing the different ATPS formulations through the microfluidic channel and observing them with a high-speed camera to assess the size of the internal phase droplets. Consistently with previous studies,[25] increasing the (i) percentage of NaCl (0 - 9 - 18 - 27 - 36 g/L) and (ii) GelMA (0 - 3 - 5 - 7 %) in the ATPS formulation resulted in an increase in the diameter of the droplets. By removing NaCl (**Figures 3a, i - ii**) and GelMA (**Figures 3c, i - ii**) from the ATPS formulation, no emulsion formation was observed, resulting in a single-phase system. In contrast, different concentrations of NaCl (**Figures 3a, iii - x**) and GelMA (**Figures 3c, iii - viii**) resulted in different sizes of GelMA droplets. Increasing the NaCl concentration, the size of the droplets varied from an average of 12.8 ± 2.6 µm for the 9 g/L NaCl formulation to an average of 52.4 ± 11.4 µm for the 36 g/L NaCl formulation (**Table 1**). Increasing the GelMA concentration also led to a progressive increase in the average droplet diameter, ranging from 14.4 ± 2.2 µm to 32.3 ± 7.4 µm at 3% w/v GelMA, respectively (**Table 2**).

**Figure 3.**
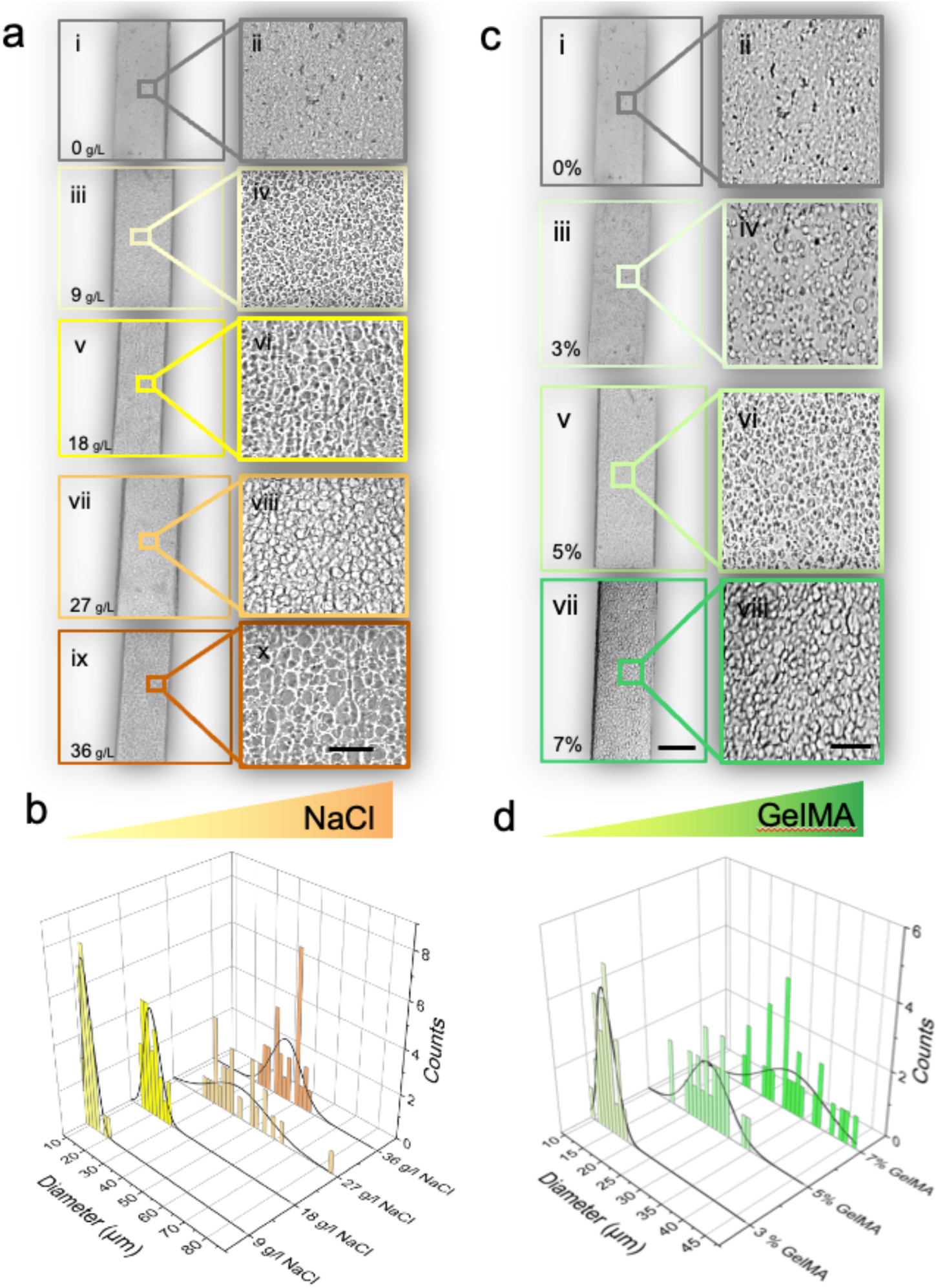
Variation of NaCl and GelMA concentrations for droplets characterisation. a) ATPS formulations differing in NaCl content visualised via FastCam. a – i, ii) N0, a – iii, iv) N9, a – v, vi) N18, a – vii, viii) N27, a – ix, x) N36. b) Droplet size distributions for different NaCl content: NaCl 9 g/L, NaCl 18 g/L, NaCl 27 g/L, NaCl 36 g/L. c) Droplet size distributions for different GelMA content: GelMA 3%, GelMA 5%, GelMA 7 %. d) ATPS formulations differing in GelMA content. d – iii, iv) G3N9, d – v, vi) N9, c – vii, viii) G7N9. Scale bars: (a – ix, c – vii) 100 μm, (a – x, c – viii) 25 μm. Mean ± S.D. n=50.

**Table 1.**
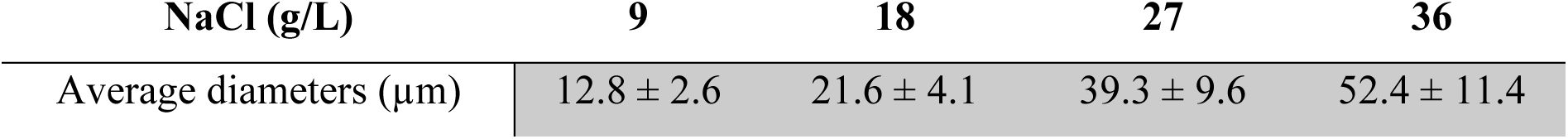
Diameter of droplets as salt concentration increases. Mean ± S.D. n=50.

**Table 2.** Diameter of droplets with NaCl (36g/L) as GelMA concentration increases. Mean α S.D. n=50.

| GelMA (% w/v) | 3 | 5 | 7 |
| --- | --- | --- | --- |
| Average diameters ($\mu\text{m}$) | $14.4 \pm 2.2$ | $21.6 \pm 4.1$ | $32.3 \pm 7.4$ |

Similarly to the effect of NaCl on the polydispersity, GelMA concentration was found able to modulate the droplet sizes of the dispersed phase in the emulsion. It is known that the increase in the average size of the droplets is due to the salting-out effect.[29] As the salt concentration increases, the ionic strength of the solution raises, thereby promoting phase separation. In this context, the reduced miscibility between the polymer components drives the formation of two immiscible aqueous phases, with GelMA preferentially segregating into the dispersed phase.[30] Considering the optimal results obtained with a GelMA concentration of 5% w/v for emulsion droplet dimensions (**Table 1**), coupled with the low-end viscosity properties ideal for microfluidic-assisted extrusion, this ATPS ink formulation was thus employed for further analysis in the present study. Indeed, elevated GelMA concentrations (e.g., 7% w/v) resulted in greater viscosity, thereby hindering flow through the microfluidic printhead and further extrusion.

A polydispersion was observed for all ATPS formulations studied (**Figures 3b, d**). Variation in the size distribution of droplets may be due to several factors, such as (i) the method used to prepare the emulsion (i.e. mechanical stirring), and (ii) the polarity of the solvent. The polarity of the NaCl + HEPES solution and its miscibility with the dispersed phase can affect the size and distribution of the droplets. Indeed, it is well established that the higher solvent polarity promotes the formation of droplets with a smaller diameter.[31] Additionally, excessive agitation of polymer solutions or prolonged heating can cause the formation of larger droplets or the breakage of droplets already formed.[32]

### 2.3 ATPs inks rheological behaviour is regulated by salt concentration and phase separation

The rheological properties of GelMA-Alginate ATPS were investigated in relation to NaCl concentration (*C_NaCl_*), revealing notable effects on viscosity, complex viscosity, and mechanical spectra. **Figure 4a** reports the shear viscosity of GelMA-Alg ATPS at a constant biopolymer concentration (5% w/v and 4% w/v, respectively) and temperature (37°C) while varying *C_NaCl_*. The relative positioning of the curves exhibits an apparently inconsistent trend. The viscosity curve at 9 g/L dominates over all the others across the entire range of shear rates investigated, decrementally followed by 36, 27, and finally 18 g/L. To provide more direct evidence of the influence of *C_NaCl_* over shear viscosity, the latter, determined at an arbitrary shear rate of 100 s^-1^, was plotted versus *C_NaCl_* (**Figure 4b**). Viscosity is at its highest value in correspondence with 9 g/l, then drops to a minimum at 18 g/L. Beyond this value, the viscosity increases approximately linearly with *C_NaCl_*. A minimum reflects the influence of two competing phenomena, each prevailing in different ranges of *C_NaCl_*. Such phenomena are the biopolymer partitioning between the two phases (particularly GelMA) and the degree of droplet packing. At 9 g/L, a relatively small amount of GelMA phase separates, forming the droplet phase. Therefore, the viscosity at 9 g/L is mainly influenced by the overall biopolymer concentration in the continuous phase, with a lesser impact from the droplet phase. The latter, characterised by a comparatively low dispersed area fraction (0.37, **Figure 2e**), are loosely packed and only interact with each other sporadically during shearing. When *C_NaCl_* reaches 18 g/l, the total amount of polymer in the continuous phase decreases because more GelMA phase separates. At the same time, the volume fraction of droplets increases to 0.61, which is just below the maximum packing degree of polydispersed droplets (0.64).[33] Evidently, at this *C_NaCl,_* the main parameter influencing viscosity is still the total polymer concentration in the continuous phase. Increasing *C_NaCl_* to 27 g/l results in a droplet fraction of 0.67 (**Figure 2e**) above the close packing limit of 0.64. ATPS viscosity is primarily influenced by the interaction and deformation of droplets during shear, with the continuous phase viscosity playing a minor role due to its lower GelMA content. The same qualitative considerations apply to 36g/L condition. The volume fraction of the droplet phase increases to 0.75 (**Figure 3e**), and as evidenced by **Figure 2a-x**, droplets are tightly packed, resulting in an overall further increase in viscosity.

**Figure 4.**
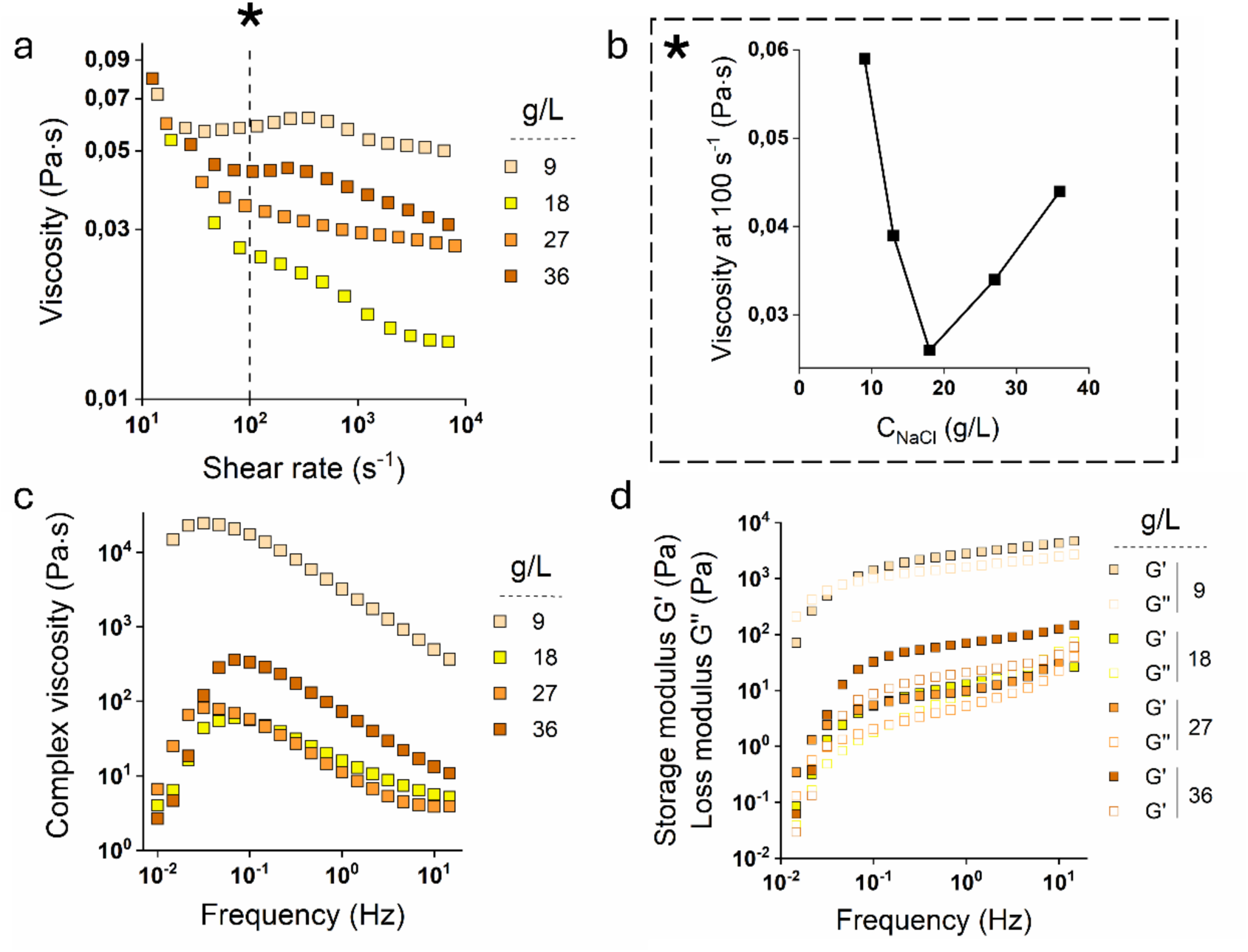
Rheological properties of GelMA-Alginate ATPS. a) Viscosity (Pa s) over shear rate (s^-1^) at increasing NaCl concentrations (% w/v); b) Viscosity at 100 s^-1^ plotted against NaCl concentration (*C_NaCl_*); c) Complex viscosity at increasing NaCl concentrations at increasing frequencies (Hz); d) ATPS mechanical spectra: storage (G’) and loss (G’’) moduli over frequency, at increasing NaCl concentrations. Mean ± S.D. n=3.

The two-phase nature of ATPS is further evidenced by the behaviour of complex viscosity (*11\**) vs frequency. Traces, as illustrated in **Figure 4c**, exhibit a peak at low frequencies (long-time scales). As the concentration of *C_NaCl_* increases, the maximum of the peaks becomes more prominent. In oscillatory experiments, part of the input energy is stored as elastic deformations, and part is dissipated. Complex viscosity is a measure of the dissipated energy. Thus, the presence of a peak indicates that at low frequencies (on the left side of the maximum), the ATPS display a more elastic behaviour. Under oscillatory stress, the interface between the two phases deforms, assuming an enhanced anisotropic shape with the interfacial area becoming larger.

Capillary pressure opposes deformation. As a result, the interface acts somewhat similarly to an elastic membrane. The absence of an evident maximum for 9g/L indicates that the interface is relatively rigid and does not deform significantly during an oscillation period. This is consistent with the smaller dimension of dispersed phase droplets (**Figure 3a** and **Table 1**), corresponding to higher capillary pressures. Additionally, the number of droplets per unit volume is smaller. As the dispersed phase volume fraction increases with *C_NaCl_*, both the droplet dimension and degree of packing increase, and the amount of energy stored increases, leading to higher peaks. As the frequency of oscillation increases, complex viscosity increases as well, indicating that deformation is increasingly out of phase with the imposed stress, and other dissipation modes such as macromolecules’ reptation take over. Such behaviour has been observed with immiscible polymer blends in the molten state.

The mechanical spectra in **Figure 4d** validate the conclusions derived from the examination of the complex viscosity curves (**Figure 4c**). The storage modulus exceeds the loss modulus in a large part of the frequency range explored, and the crossover frequency (*f_c_*) shifts to the higher frequency side (shorter relaxation times) as the NaCl content increases, indicating enhanced elastic properties. Therefore, as the degree of droplet packing increases, the ATPS acquire increasingly pronounced solid-like characteristics.[34]

### 2.4 Selective removal of ATPS phases can modulate degradation and material release

The production of ATPS fibres can be tailored by manipulating the NaCl content to yield either monophasic or biphasic systems. In monophasic systems (e.g., 0 g/L NaCl), phase separation does not occur, and alginate and GelMA are homogeneously mixed within the fibre. Following cross-linking, GelMA functions as a structural binder, thereby contributing to the mechanical integrity of the fibre, even in the absence of alginate. Conversely, in biphasic systems formed at higher NaCl concentrations (e.g., 36 g/L), phase separation promotes the segregation of GelMA into the internal dispersed phase, with alginate forming the continuous outer matrix. To confirm the distribution of the two polymer phases and their respective roles in maintaining scaffold integrity, a selective dissolution of both alginate and GelMA was performed. In the selective removal of alginate, the behaviour of the 3D constructs (cross-linked both chemically and physically) was observed at 0 g/L NaCl and 36 g/L NaCl before and after the use of the enzyme alginate lyase (**Figure 5a**).

**Figure 5.**
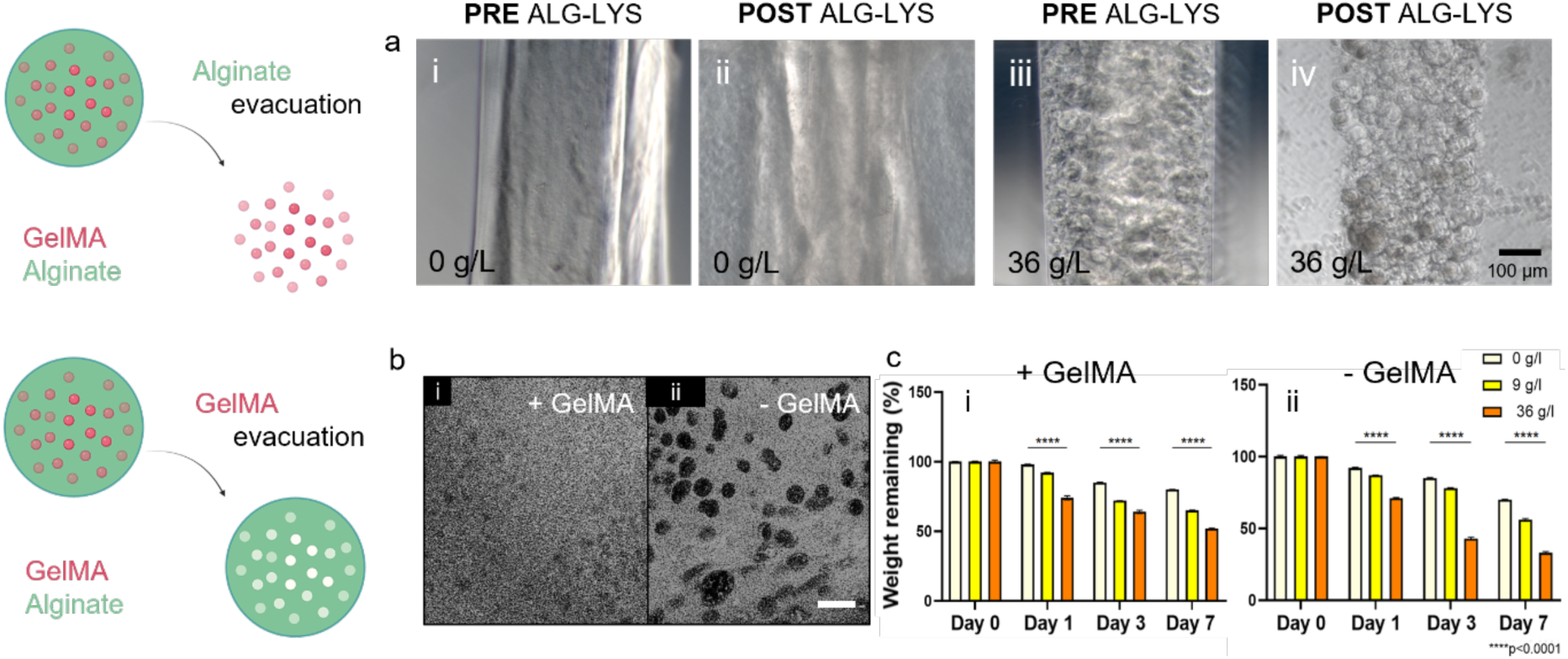
Selective degradation of GelMA and alginate in ATPS systems. (a) Selective degradation of alginate into ATPS fibres. Monophasic samples at 0 g/L NaCl a-i) before and a-ii) after incubation with alginate lyase. Biphasic samples at 36 g/L NaCl a-iii) before and a-iv) after alginate lyase. b) Selective degradation of GelMA in ATPS scaffolds. Confocal images with b-i) GelMA maintenance and with b-ii) GelMA removal. c) Degradation test performed on samples at 0 – 9 – 36 g/L. Degradation test on samples in which c-i) GelMA was chemically cross-linked and alginate was physically reticulated and degradation test on samples in which c-ii) GelMA was not chemically cross-linked, and alginate was physically reticulated. Scale bars: (a) 100 μm, (b) 25 μm. Statistical significances were assessed by one-way ANOVA. Mean ± S.D. n=3, ****p<0.0001.

In the case of the monophasic fibres (0 g/L NaCl), partial degradation was observed (**Figures 5a, i – ii**), but the fibre structure remained largely intact. This phenomenon can be attributed to the homogeneous distribution of GelMA, which provides internal cohesion and stability once cross-linked, independently of the alginate. In contrast, biphasic fibres produced with 36 g/L NaCl displayed complete structural collapse upon enzymatic degradation of alginate (**Figures 5a, iii – iv**), causing the release of GelMA droplets. This observation is consistent with the phase-separated structure of the scaffold, in which GelMA is mostly confined to internal domains while alginate forms the continuous external matrix. When alginate is removed, the external phase that binds the internal GelMA-rich regions collapses, resulting in the degradation of the entire construct. These findings support the structural role of alginate as a matrix-forming component in biphasic systems and the lack of cohesion between isolated GelMA domains when alginate is removed.

Selective removal of GelMA was also performed to assess the impact of internal GelMA domains on scaffold architecture. Scaffolds with 9 g/L salt were produced by physically cross-linking alginate only. The samples were placed in an incubator at 37 °C for 2 h to eliminate the chemically non-cross-linked GelMA. As a control, equal samples were prepared but stored in a 4 °C fridge after the physical cross-linking of the alginate to prevent the GelMA from dissolving and consequently being removed. Following GelMA removal, the fibre structure exhibited a discontinuous morphology, with voids corresponding to the previously GelMA-rich regions. In contrast, the control samples maintained the biphasic emulsion-like appearance (**Figures 5b, i – ii**).

A 7-day *in-vitro* degradation test was further performed to assess the stability of the scaffolds and their ability to degrade over time with both cross-linked and chemically non-cross-linked GelMA as an internal emulsion phase. It was observed that the samples in which GelMA was not chemically cross-linked degraded more rapidly compared to the structures in which GelMA was cross-linked and maintained within the scaffolds (**Figures 5c, i – ii**). The result revealed a higher stability of the cross-linked structures, thus highlighting how the cross-linking process significantly (p<0.0001) alters the mechanical properties and resistance to degradation of the scaffolds. The formulation that includes cross-linked GelMA tends to be more resistant to hydrolytic degradation due to the packed covalent junctions within the GelMA network. On the other hand, as confirmed in the work of Velasco and co-workers,[35] non-cross-linked GelMA degrades rapidly under physiological conditions, as lacks the stability provided by the cross-linking network.

### 2.5 Microfluidic-assisted 3D bioprinting of ATPSs facilitates the patterning of biphasic structures

The rheological analysis revealed that ATPS, independently of the NaCl content, do not possess the level of viscoelasticity (of the order of hundreds or thousands Pa⸱sec^-1^) necessary for their direct printing. Thus, we have here employed a microfluidic-assisted 3D bioprinting approach to effectively deposit low-viscosity bioinks. In this work, we used a custom-made flow-focusing microfluidic printhead. In this configuration, the ATPS flows in the central microchannel (core flow) while the cross-linking CaCl_2_ solution is delivered through the lateral channels (sheath flow), using a previously reported monolithic PDMS printhead.[22,23] The CaCl_2_ solution drives the instantaneous ionotropic gelation of the alginate present prevalently in the ATPS continuous phase, facilitating the deposition of biphasic and monophasic lattice structures in-air layer by layer (**Figure 6a**).

**Figure 6.**
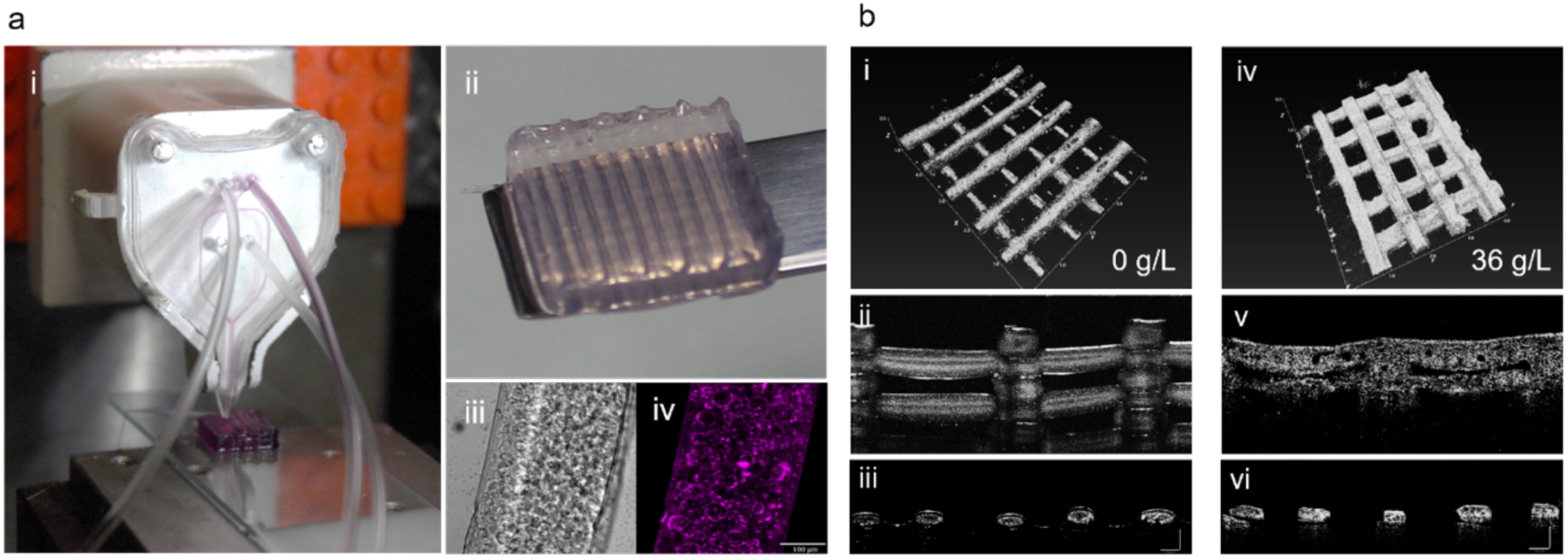
Microfluidic 3D bioprinting of ATPSs. a-i) Image of the microfluidic printhead during 3D deposition of an ATPS scaffold. a-ii) Air-printed ATPS scaffold lattice layer by layer with detail of a single fibre in a-iii) brightfield and in a-iv) alginate autofluorescence at 405 nm. b) OCT images of the ATPS systems. b-i) Printed grid image for the 0 g/L NaCl composition, b-ii) frontal fibre image, b-iii) fibre sections. b-iv) Printed grid image for the 36 g/L NaCl composition, b-v) frontal fibre image, b-vi) fibre sections. Scale bar: (a, iii – iv) 100 µm. Mean ± S.D. n=3.

To investigate the internal structure of the printed scaffolds in the wet state, an OCT analysis was performed (**Figure 6b**). After producing the scaffolds without chemical cross-linking of GelMA, the latter was dissolved overnight in an incubator for the complete removal of the internal phase of the scaffolds. As confirmed in previous analyses, the results showed that the monophasic 0 g/L salt scaffolds had a compact and homogenous internal structure, whereas the biphasic 36 g/L NaCl scaffolds were characterised by internal apparent pores.

The ATPS fibres were also evaluated for swelling up to 24 h after their production, and a non-significant decrease in fibre diameter was observed after 12 h, followed by an increase in fibre size for both monophasic (0 g/L NaCl) and biphasic (36 g/L NaCl) systems (**Figures S1a-b**). Importantly, the fibres were found to consist of two polymer-rich immiscible compartments, namely alginate-rich and GelMA-rich phases. The slight discrepancies observed in swelling behaviour can be due to disparities in ionic strength. Specifically, the enhanced swelling exhibited by ATPS fibres with 36 g/L NaCl can be ascribed to the gradual release of NaCl into the surrounding medium. This release might reduce the ionic shielding of the negatively charged alginate chains, allowing for greater electrostatic repulsion and subsequent expansion of the alginate rich-phase.[36]

Ultimately, mechanical tests were also carried out to evaluate the storage and loss modulus of 3D scaffolds in which both cross-linked and non-cross-linked GelMA were studied (**Figure S1c**). All hydrogel samples containing GelMA showed distinction between *E’* and *E’’* values in the 100-200 kPa and 20-100 kPa range respectively, confirming the predominantly elastic response with greater stiffness values registered for the composition at 9 g/L NaCl content. This parallels the rheological results, as at this salt content, the majority of the polymer is concentrated in the continuous phase. Elastic and characteristic gel behaviour was also observed in the case of GelMA-free compositions at a concentration of 0 g/L and 9 g/L NaCl. In details, the slight increase in modulus observed for the composition containing 9 g/L salt following the removal of GelMA may be related to a more homogeneous distribution of the alginate network under those ionic conditions. In contrast, for the composition containing 36 g/L NaCl, a divergent mechanical behaviour was observed following GelMA removal, manifesting as a predominantly viscous response. This is potentially attributable to the presence of non-cross-linked GelMA within the internal structure of the material.

Indeed, scaffolds formed at 36 g/L salt showed the presence of internal voids resulting from the removal of the segregated GelMA-rich phase. In addition, SEM analysis was conducted to assess the internal architecture of the scaffolds. From the acquired images, it was possible to confirm that the fibres of the monophasic system (0 g/L NaCl) were found homogenous and smooth in surface, while the ATPS structure (36 g/L NaCl) showed evident internal interconnected porosity (**Figure S2**). Importantly, the porosity should not be interpreted as derived from an intrinsically porous structure, but rather as phase-induced domains created by the biphasic nature of the ATPS formulation.

### 2.6 Cells remain viable following extrusion and preferentially spread over the dispersed phase with higher NaCl concentration

To evaluate the cytocompatibility of the developed ATPS scaffolds and to investigate how the internal architecture and phase separation influence the distribution and morphology of the cells, C2C12 murine myoblasts were used as a model. These cells are recognised for their sensitivity to biochemical and mechanical signals from the surrounding matrix,[37] thus serving as suitable model to assess the bio-instructive potential of ATPS-derived scaffolds.

C2C12 cells were encapsulated in ATPS inks at increasing NaCl concentrations to observe whether phase separation differences, determined by salt concentration, could drive the preferential localisation and diffusion of cells, and to determine how the presence or removal of the GelMA-rich phase influenced these behaviours. Elevated viability was observed in all conditions up to 7 days of scaffold culture, with an increase at day 7 compared to day 1 (**Figures 7a, i**), confirming the non-toxic effect of NaCl inclusion.

**Figure 7.**
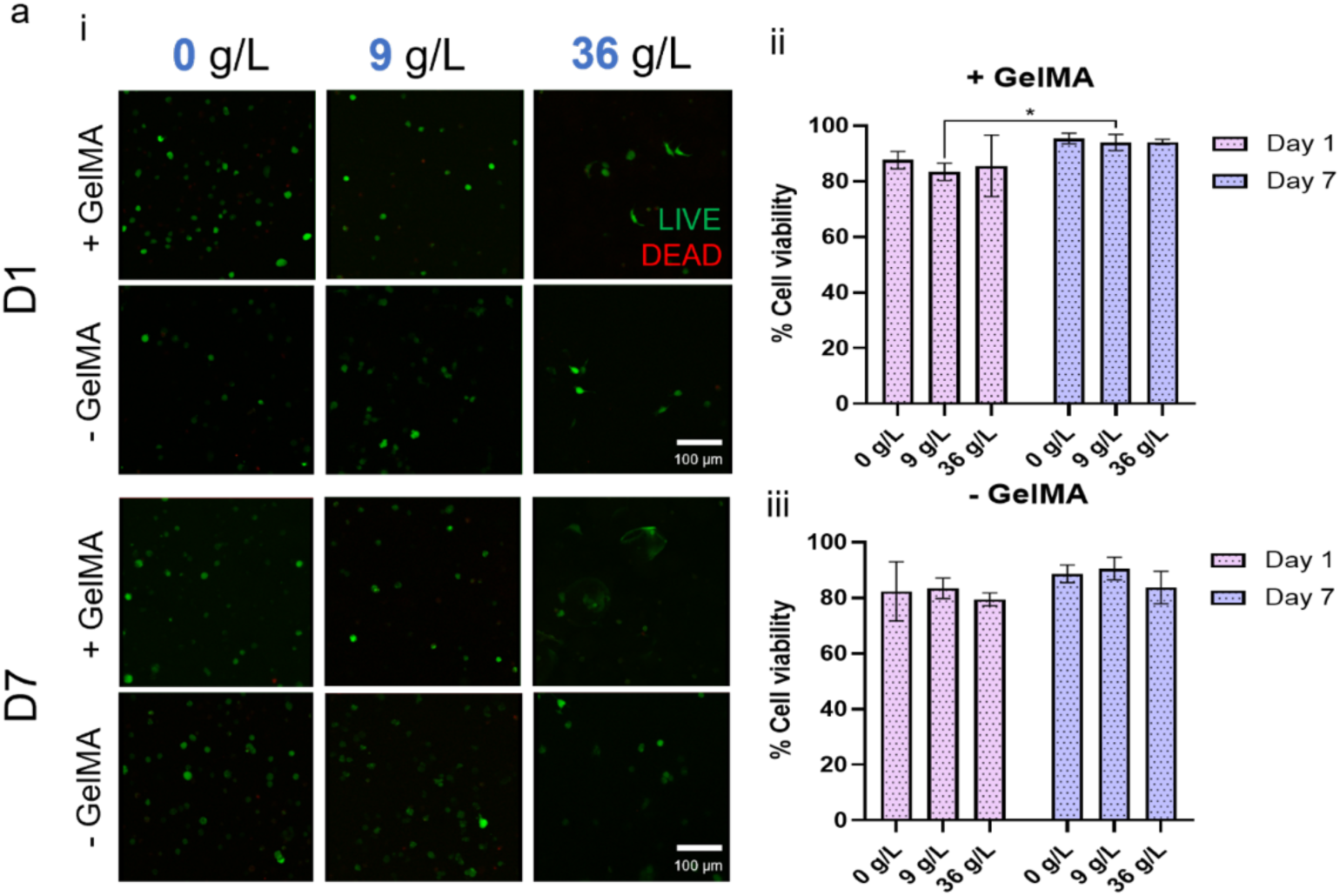
Cell viability of C2C12 in ATPS systems. a-i) Confocal images with C2C12 labelled with Calcein to show live cells (green) and cells labelled with propidium iodide to show dead cells (red). Quantification of cell viability in ATPS systems where a-ii) GelMA was cross-linked and in samples where a-iii) GelMA was removed. Scale bars: (a) 100 µm. Statistical significances were assessed by two-way ANOVA. Mean ± S.D. n=3, *p<0.05.

Viability was found to increase significantly (p<0.01) over 7 days of culture (**Figures 7a, ii**) in the presence of GelMA. However, following GelMA removal (**Figures 7a, iii**) the viability was found non-significantly consistent over the period of 7 days. Studies have shown that cells encapsulated in GelMA exhibit high viability, promoting cell adhesion and proliferation over time.[38] In contrast, the use of alginate as a biomaterial generally requires modifications to improve cell adhesion and viability.[39] The lack of these modifications in the molecular structure of alginate chains often results in a reduced cell viability, as was confirmed from our analyses in which alginate scaffolds were cleaned of their internal GelMA phase. By culturing 3D bioprinted and marking the nuclei and actin filaments of the cells, the spatial arrangement and spreading of C2C12 inside the printed ATPS were assessed (**Figure 8a-d**).

**Figure 8.**
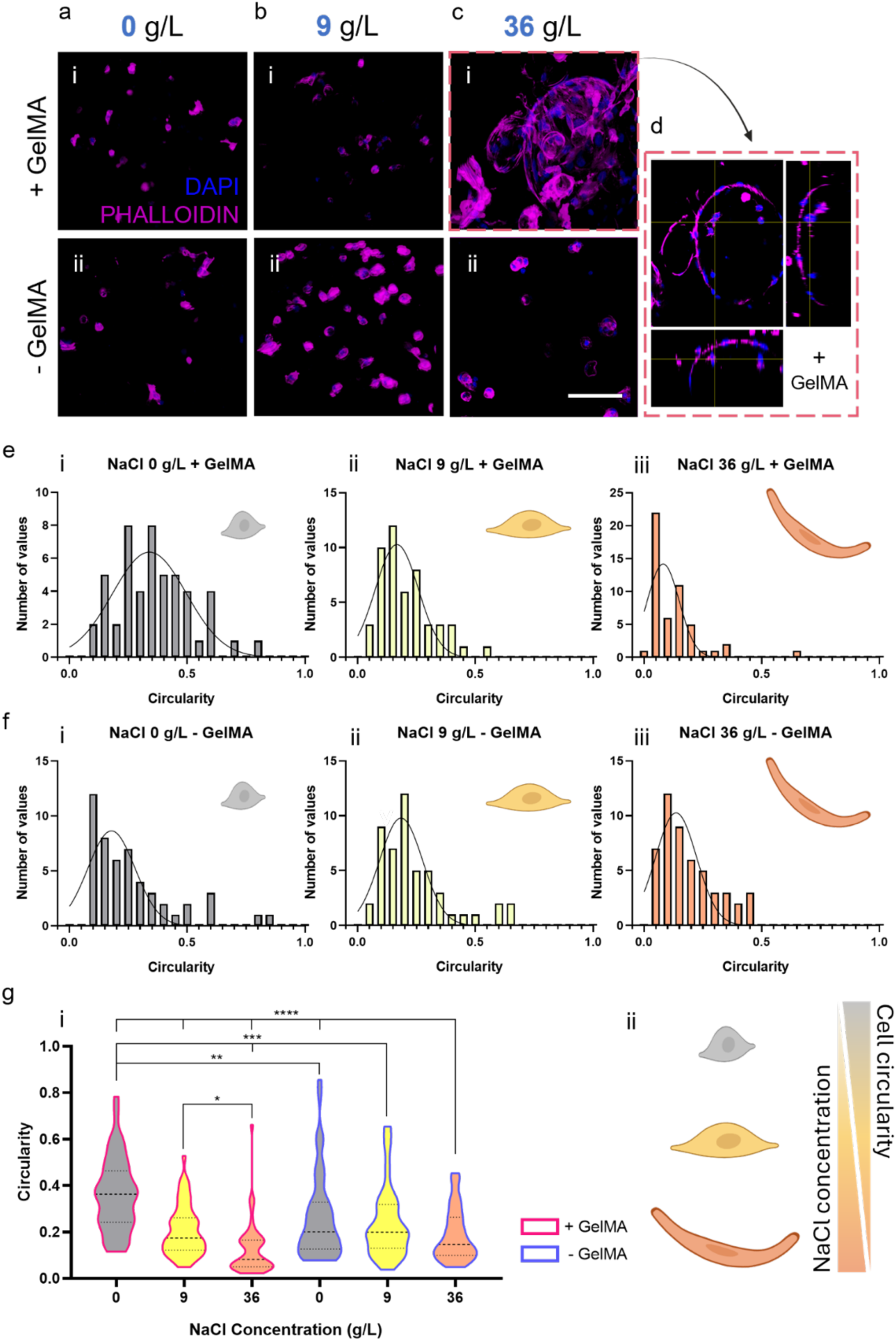
Spatial distribution of C2C12 in ATPS systems. a-b-c) Confocal images with C2C12 labelled with DAPI to show nuclei (blue) and cells marked with phalloidin to show f-actin (magenta). d) Orthogonal view of image c-i. Circularity analysis conducted on the formulation e-i) NaCl 0 g/L with GelMA, e-ii) NaCl 9 g/L with GelMA, e-iii) NaCl 36 g/L with GelMA, f-i) NaCl 0 g/L without GelMA, f-ii) NaCl 9 g/L without GelMA and f-iii) NaCl 36 g/L without GelMA. g) Cell circularity as a function of salt concentration. g-i) Cell circularity for each NaCl composition within the ATPS samples is shown. The pink contour marks samples with + GelMA; the blue contour marks samples with - GelMA. g-ii) Cell circularity is inversely proportional to the salt concentration in the hydrogels. When the amount of NaCl in the scaffolds is higher, the cells are more elongated. Scale bars: (c) 100 µm. Statistical significances were assessed by two-way ANOVA. Mean ± S.D. n=3, ****p<0.0001, ***p<0.001, **p<0.01, *p<0.05.

From confocal analysis, it can be appreciated that in the absence of GelMA as ATPS internal phase, C2C12 were arranged in a relatively random pattern, with no particular spreading within the ATPS. In contrast, 3D bioprinted structures with chemically cross-linked GelMA, cells in the 0 - 9 g/L systems presented a similar spreading behaviour, with the C2C12 in the 9 g/L NaCl material being more located in the GelMA droplets. The distribution of C2C12 within the 36 g/L NaCl material, on the other hand, was markedly more oriented inside the inner phase droplets, where the cells were concentrated inside and grew along the entire surface of the GelMA droplets. This behaviour is attributable to the enhanced cell adhesion properties of GelMA compared to alginate.[40] The RGD (Arg-Gly-Asp) sequences of GelMA favour cell organisation, creating an interconnected network in the inner phase droplets.[41] The alginate phase in the ATPS formulation, favour the reduction of cell spreading, demonstrating a larger density of rounded and isolated cells.[42] The increased salt concentration in ATPS results in higher phase separation between GelMA and alginate, causing cells to localise more in GelMA-rich regions. Circularity analyses performed on C2C12 within the ATPS hydrogels confirmed what was observed by confocal analysis (**Figures 8e-g**). In fact, the elongated cells in the 36 g/L systems showed significant circularity values tending closer to 0 (0: non-circular - 1: circular), whereas the 0 - 9 g/L cells were more round with higher circularity values.

To assess whether the spatial arrangement of C2C12 within the ATPS-based biomaterial was a standard behaviour, DAPI was used to label the nuclei of various cell types (A549, C2C12, MG63, HBMSCs) encapsulated in ATPS inks in which both emulsion components were cross-linked, and in samples in which cross-linking was carried out exclusively for alginate, favouring GelMA evacuation (**Figure S3**).

The latter condition was essential to identify, depending on the salt concentration used (0 - 9 - 36 g/L NaCl), in which of the two phases (outer phase of alginate or inner phase of GelMA) the cells were located. The results showed that in a monophasic system (0 g/L NaCl), cells arranged randomly and non-homogeneously. In contrast, ATPS (9 - 36 g/L NaCl) supported the organisation of most of the cells surrounding as well as within the GelMA-rich droplets and to a lesser extent in the outer alginate-rich phase. In particular, cells were found increasingly stimulated to arrange themselves within the dispersed phase, when encapsulated in ATPS with 36 g/L NaCl, thus larger GelMA-prevalent droplets. This behaviour aligned with previous research, suggesting that while GelMA enhanced cell adhesion and organization due to its RGD motifs,[29] alginate offered excellent structural support and stability,[43] which can be further leveraged to modulate cell behaviour. Indeed, alginate facilitated the formation of well-defined microenvironments that promoted cellular encapsulation, protection, and growth, particularly in high-salt conditions where phase separation promotes localised cell clustering.

### 2.7 HBMSCs spreading ratio increases with higher salt dispersion in 3D printed constructs

HBMSCs are a promising cell source for skeletal TE applications due to their inherent osteogenic potential and ability to respond to microenvironmental cues.[44] In this context, the functionality of HBMSCs in ATPS structures was investigated, both in terms of cell spreading/viability and the osteogenic differentiation capacity of the cells depending on the different salt concentrations. Following the observation of more consistent behaviour in conditions where GelMA was preserved and cross-linked within ATPS hydrogels, the experiments were narrowed to only those formulations containing cross-linked GelMA (0, 9, and 36 g/L salt). Consequently, experiments on formulations without cross-linked GelMA were no longer performed. In the selected scaffolds, alginate was cross-linked physically, while GelMA was cross-linked chemically. The viability of the HBMSCs within ATPS was evaluated at increasing concentration of NaCl, noticing a non-significant change in the percentage of viable stromal cells up 7 days of culture, confirming the biocompatible nature of ATPS even at the highest salt concentration (**Figure S4**).

Although alginate is recognised for its limited cell-adhesion properties,[45] the incorporation of GelMA in the ATPS formulations enabled the overcoming of this limitation, thereby supporting cell viability across all tested conditions. As outlined above, the GelMA and alginate-based ATPS synergistically exploited the complementary properties of both biomaterials: GelMA provided high cell adhesion and promotes cell,[29] while alginate provided superior structural integrity and mechanical stability.[43] This collaboration resulted in improved cell behaviour and functionality within the composite ATPS hydrogels. The cellular distribution of HBMSCs in ATPS hydrogels (**Figure 9a**) was then confirmed and the spreading assessed by staining of nuclei and f-actin (**Figure 9b**). HBMSCs were found to preferentially concentrate within the GelMA phase of 36 g/L salt samples, colonizing the entire volume and preferentially growing around the phase boundaries.

**Figure 9.**
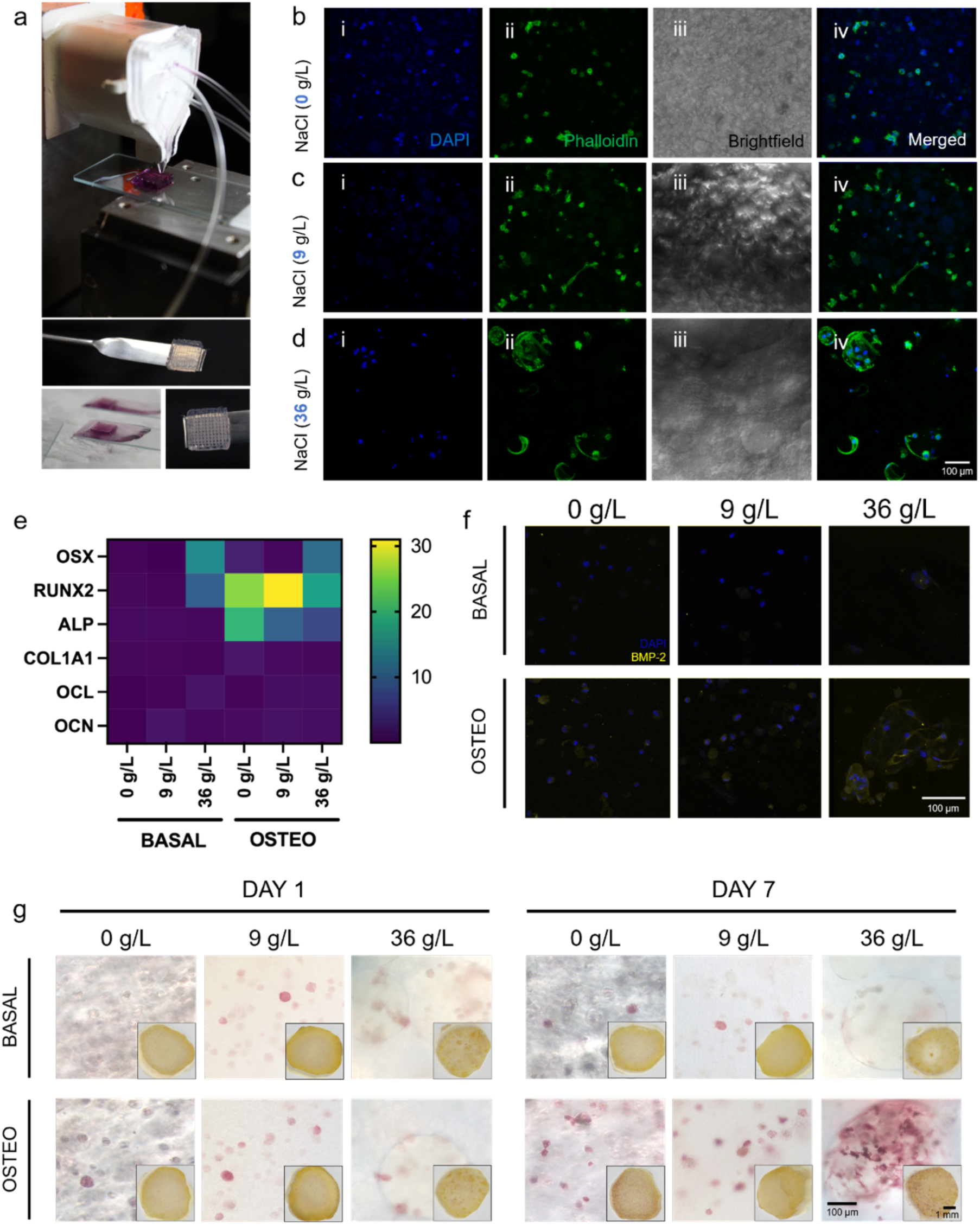
Spreading and osteogenic differentiation of HBMSCs in ATPS hydrogels at different salt concentrations. a) Microfluidic 3D bioprinting of HBMSCs ATPS-scaffolds. b-c-d) Confocal images with HBMSCs labelled with DAPI to show nuclei (blue) and cells marked with phalloidin to show f-actin (green). e) RT-qPCR analysis of HBMSCs after 7 days of 3D culture in basal and osteogenic media. The protein expression of further factor associated with osteogenic differentiation was examined to observe the effect of different culture media on ATPS scaffolds. The expression levels of osterix (OSX), RUNX2, alkaline phosphatase (ALP), COL1A1, osteonectin (OCL) and osteocalcin (OCN) were found higher in ATPS constructs cultured in osteogenic media at day 7 compared to controls in basal media. f) Immunofluorescence images with HBMSCs labelled with DAPI to show nuclei (blue) and cells marked with BMP-2 monoclonal antibody to show BMP-2 (yellow). g) Microscopic and macroscopic (insert) view of alkaline phosphatase staining performed at day 1 and day 7 in samples at 0 – 9 – 36 g/L NaCl treated in basal and osteogenic medium. Scale bars: (b, f) 100 µm, (g) 100 µm – 1 mm. Statistical significances were assessed by two-way ANOVA. Mean ± S.D. n=3.

To assess the capacity for osteogenic differentiation, the expression of specific genes of HBMSCs encapsulated in ATPS hydrogels was determined by RT-qPCR (**Figure 9e**). ATPS scaffolds were maintained in culture with both basal and osteogenic medium. A pronounced increase was observed at day 7 in the levels of the OSX, RUNX2 and ALP genes when the scaffolds were cultured in osteogenic medium. The expression of a further gene related to osteogenic differentiation (BMP-2) was observed by confocal microscopy in ATPS scaffolds at increasing salt concentrations (**Figure 9f**). Higher expression was detected in the case of the hydrogels at 36 g/L salt when they were cultured in osteogenic medium. In particular, the GelMA droplets appeared to stimulate and enhance the accumulation of BMP-2 within the dispersed phase. The increase in gene expression suggests that the presence of GelMA provides an ideal microenvironment for osteogenic differentiation. Indeed, the presence of BMP-2 demonstrates bone matrix production and mineralisation[46] in ATPS scaffolds. Finally, alkaline phosphatase (ALP) staining revealed a substantial difference between ATPS scaffolds treated in basal medium and osteogenic medium up to 7 days (**Figure 9g**). In particular, at day 1 there was no particular difference in the level of ALP expression, whereas at day 7 the difference between samples in basal medium and samples in osteogenic medium was marked. Indeed, when treated in osteogenic medium, ALP expression was notably higher, with greater disposition and expression localised within the GelMA droplets. This is consistent with the expected timing of osteogenesis, in which ALP expression increases as cells start to deposit the initial bone matrix.[47] The observation of the higher ALP expression in the GelMA droplets further underlines the role of GelMA as a bioactive component that increases the osteogenic potential of the encapsulated HBMSCs.

### 2.8 Evaluation of VEGF-mediated vascular ingrowth and AI-assisted quantification using CAM models

To assess the potential for tissue repair and vessel regrowth, six fertilised eggs per experimental condition were used to host the ATPS bioconstructs and implanted for one week.[48] VEGF-mediated vessel growth showed a significant increase compared to the control (-VEGF) groups after seven days of incubation (**Figure 10a**). In particular, the CAM model showed increased vascularisation in scaffolds with higher salt concentration (36g/L) (**Figure 10a, iii-iv**). The constructs at 0 g/L maintained their structural integrity for a period of 24 h following implantation and at the time of isolation, suggesting minimal integration with the surrounding environment. In contrast, the 36g/L constructs demonstrate evident signs of loosening and integration within the CAM model 24 h after implantation (**Figure 10b**). Quantitative analysis revealed a greater number of afferent vessels in the 36g/L+VEGF groups compared to the -VEGF groups (**Figure 10c**). Notably, vessel density in the 36g/L models increased two folds in the +VEGF groups compared to the 0 g/L and – VEGF control groups, indicating a strong angiogenic response. The results suggest that the 36g/L models with VEGF stimulated new vessel formation, whereas the vessels in the other groups remained immature and unevenly distributed.

**Figure 10.**
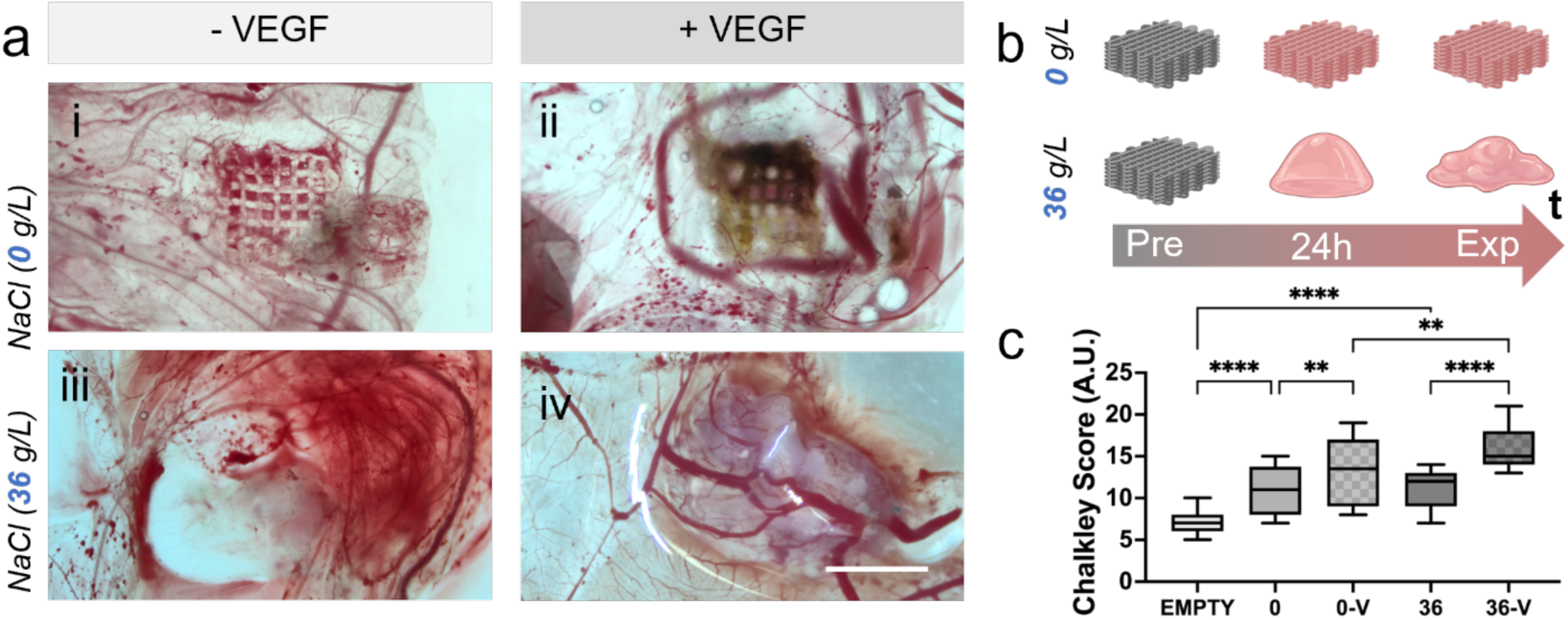
Evaluation of vascularisation of ATPS constructs in a CAM model. a) Macrographs of the explanted constructs of (i,ii) 0 g/L and (iii, iv) 36 g/L, without or with VEGF, respectively. b) Rendering of the dynamic changes in architectures observed comparing the fabricated constructs (Pre) and the resulting explanted platforms (Exp). c) Chalkley score of the vascularised implants. Scale bars: (b) 10 mm. Mean ± S.D, n=6, *p<0.05, **p<0.01, ****p<0.0001.

The present results are consistent with previous studies that have demonstrated the pro-angiogenic potential of localised VEGF compounds.[49] At the same time, the increased salt concentration and the compartmentalisation of GelMA might have played a central role in fostering an augmented angiogenic response. Indeed, previous findings have also demonstrated the pro-angiogenic potential of sodium compounds in CAM models. For instance, sodium nitrite has been found to significantly promote angiogenesis, thereby reinforcing the hypothesis that specific sodium environments may enhance blood vessel formation.[50] However, other reports suggest that excessive salt exposure may impair angiogenesis, resulting in reduced vascular density in the CAM model.[51] This suggests that the impact of salt on angiogenesis is highly dependent on the concentration and specific type of salt involved.

Further *in vivo* studies with longer incubation times and the incorporation of co-stimulatory factors are essential to evaluate the long-term stability, perfusion capacity and functional integration of the induced vascular network for potential regenerative applications.

## 3. Conclusion

The development of ATPS bioinks for 3D bioprinting represents a significant advancement in TERM. By finely tuning the composition of GelMA and alginate, as well as the concentration of NaCl, we successfully controlled the formation and properties of W/W emulsions. The tuneable NaCl salt concentration exerted a direct influence on the size and packing degree of the dispersed GelMA droplets within the alginate phase, thereby enabling the fabrication of well-defined and complex structures. Notably, mechanical analysis demonstrated a clear relationship between salt concentration and the mechanical properties of the scaffolds, with higher NaCl concentrations resulting in a biphasic and elastic structures, which was validated by both rheological testing and OCM. Cell encapsulation (A549, C2C12, MG63, and HBMSCs) showed that the ATPS inks supported cell viability, spreading, and differentiation over time. Most importantly, ATPS induced the agglomeration of cells in small pockets, resulting in an effective compartmentalisation inside the 3D printed fibre. In particular, HBMSCs cultured in ATPS scaffolds exhibited significant osteogenic differentiation, with enhanced expression of key markers such as RUNX2 and BMP-2 when cultured in osteogenic medium, confirming the potential to guide bone tissue regeneration. The CAM assay further validated the biocompatibility of the constructs, demonstrating significant vascularisation and scaffold integration at higher NaCl concentrations. This work demonstrated, for the first time, that ATPS-based hydrogels, which are typically difficult to print due to their low-end rheological properties, can be successfully processed using microfluidic-assisted 3D bioprinting. In contrast to current approaches relying on light-based techniques, this study showed that microfluidic-assisted 3D bioprinting technology enabled the stable fabrication of ATPS scaffolds with a well-defined internal architecture, highlighting its critical role in maintaining phase separation and ensuring controlled mixing during printing, while promoting cellular functionality. Altogether, the versatility of ATPS biomaterial inks offers a powerful tool for engineering complex tissue architectures with microscale precision. The ability to selectively control cell behaviour and scaffold architecture provides a unique opportunity for creating functional tissue substitutes, paving the way for future clinical applications in regenerative medicine.

## 4. Materials and methods

### Materials

Gelatin from porcine skin (∼300 g Bloom, Type A), Dulbecco′s Phosphate Buffered Saline (PBS), Hanks′ Balanced Salt solution (HBSS), 4-(2-Hydroxyethyl)piperazine-1-ethanesulfonic acid (HEPES), Methacrylic anhydride (MA), N-Hydroxysuccinimide (NHS), N-hydroxysulfosuccinimide sodium (sulfo-NHS), N-(3-Dimethylaminopropyl)-N′-ethylcarbodiimide hydrochloride (EDC), Dimethylformamide (DMF), Sodium chloride (NaCl), Calcium chloride (CaCl_2_), Glycerol, Rhodamine B, Fluorescein 5(6)-isothiocyanate (FITC), Tris(2,2’-bipyridyl)dichloro-ruthenium(II) hexahydrate (Ru), Sodium persulfate (SPS), Alginate Lyase, Collagenase D, Dulbecco′s Modified Eagle′s Medium – high glucose (DMEM), Minimum Essential Medium Eagle – Alpha modification (αMEM), L-Glutamine, Penicillin-Streptomycin, Fetal bovine serum – non-USA origin (FBS), Trypsin-EDTA solution (1x), β-Glycerophosphate disodium salt hydrate (β-GP), 2-Phospho-L-ascorbic acid trisodium salt (AA2P), Dexamethasone, Formaldehyde solution (37 % in water), Triton™ X-100, Bovine Serum Albumin (BSA), 4′,6-Diamidino-2-phenylindole dihydrochloride (DAPI), Naphtol AS-MX Phosphate Alkaline, Fast Violet Salt, 3,4-Dihydroxy-9,10-dioxo-2-anthracenesulfonic acid sodium salt (Alizarin Red S), Ammonium hydroxide solution (25%), Lysozime and Protein Quantification Kit-Rapid were purchased from Sigma Aldrich. Calcein (AM, cell-permeant dye), Propidium Iodide, Goat anti-Rabbit IgG (H+L) Cross-Adsorbed Secondary Antibody – Alexa Fluor™ 488, Goat anti-Mouse IgG (H+L) Highly Cross-Adsorbed Secondary Antibody – Alexa Fluor™ 594, RUNX2 Recombinant Polyclonal Antibody, BMP-2 Monoclonal Antibody, Osteopontin Monoclonal Antibody, Alexa Fluor™ 488 Phalloidin, CellPath OCT Embedding Matrix and TRIzol Reagent were obtained by ThermoFisher Scientific. Alginate (Protanal® GP 1740), Ethanol absolute anhydrous (EtOH), SYLGARD™ 184 Silicone Elastomer Kit (PDMS) and Invicta 907 was purchased from FMC BioPolymer, Carlo Erba (Italy), Dow and Rimas Engineering (Italy) systems, respectively. Recombinant Human Vascular Endothelial Growth Factor-A (rh VEGF-A) was obtained by ImmunoTools (Germany).

### Synthesis of Gelatin Methacryloyl

GelMA was produced following a procedure published elsewhere.[52] Gelatin was dissolved at 10% w/v in PBS at a controlled temperature of 50 °C. After 1 h, MA (0.8 g per gram gelatin) was added, and the reaction stopped after 3 h. The solution was then transferred to 1 kDa cut-off dialysis tubes and the deionised water (DW) was changed twice daily for 5 days. Finally, the GelMA obtained was frozen (−80 °C), lyophilised (LyoQuest, Teslar, Italy) and stored in the fridge at 4 °C until further use.

### Synthesis of Gelatin Methacryloyl-Rhodamine B

GelMA-Rhodamine B was synthesised as previously described.[53] Briefly, 1 g of GelMA was dissolved in 20 ml of DW at 50 °C and reacted with 1 ml of activating solution for 24 h in the dark. The activating solution was prepared by dissolving 0.01g Rhodamine B, 0.012g NHS and 0.02g EDC in 1 ml DMF and keeping it stirring for 4 h at room temperature (RT). The solution was then purified via dialysis (1 kDa cut-off, 5 days), freeze-dried and stored in the refrigerator at 4 °C in the dark until use. *Synthesis of Alginate-FITC*: Alginate-FITC was prepared according to a previous protocol.[54] Briefly, 0.96 g of purified alginate was dissolved in 150 ml of PBS (pH 7.5). Subsequently, 0.054 g EDC and 0.06 g sulfo-NHS were added to the solution and left to stir for 2 h. Following, 0.09 g FITC was added and the reaction was left stirring in the dark for 24 h. The product was purified via extensive dialysis (kDa cut-off) using sequential washes in DW and NaCl (1 M), then freeze-dried and finally stored in the dark at 4 °C until required.

### Fabrication of microfluidic printhead

The microfluidic printing head was manufactured in PDMS with the following characteristic dimensions: 100 µm biomaterial phase channel; 200 µm CaCl_2_ and glycerol channel; 300 µm output channel. The fabrication followed a procedure reported previously.[22] Briefly, a microfluidic printhead mould was designed using a CAD software (Autodesk Fusion 360) and printed with a SLA printer (XFAB 3500SD, DWS, Italy). The mould was washed extensively in EtOH, UV-cured (MeccatroniCore BB Cure) and stored at 80 °C overnight. PDMS was then cast in the mould and polymerised at 70 °C. The cured PDMS was then bonded to the other half by plasma treatment for 2 minutes.

### ATPS preparation

ATPS materials were produced by mixing different concentrations of GelMA (3 - 5 - 7 %), alginate (4 %), Ru (1 mM) and SPS (10 mM) in an aqueous solution based on different concentrations of NaCl (0 - 9 - 18 - 27 - 36 g/L) and HEPES (25 mM) (**Table 3**). For fluorescent ATPSs, GelMA-Rhodamine B and alginate-FITC were used. The solution was left to stir at 300 RPM for 3 h, with Ru/SPS added just before printing to initiate GelMA cross-linking. The emulsion ink was then flowed into the microfluidic chip via the control of microfluidic pumps (neMESYS, Cetoni GmbH).

**Table 3.**
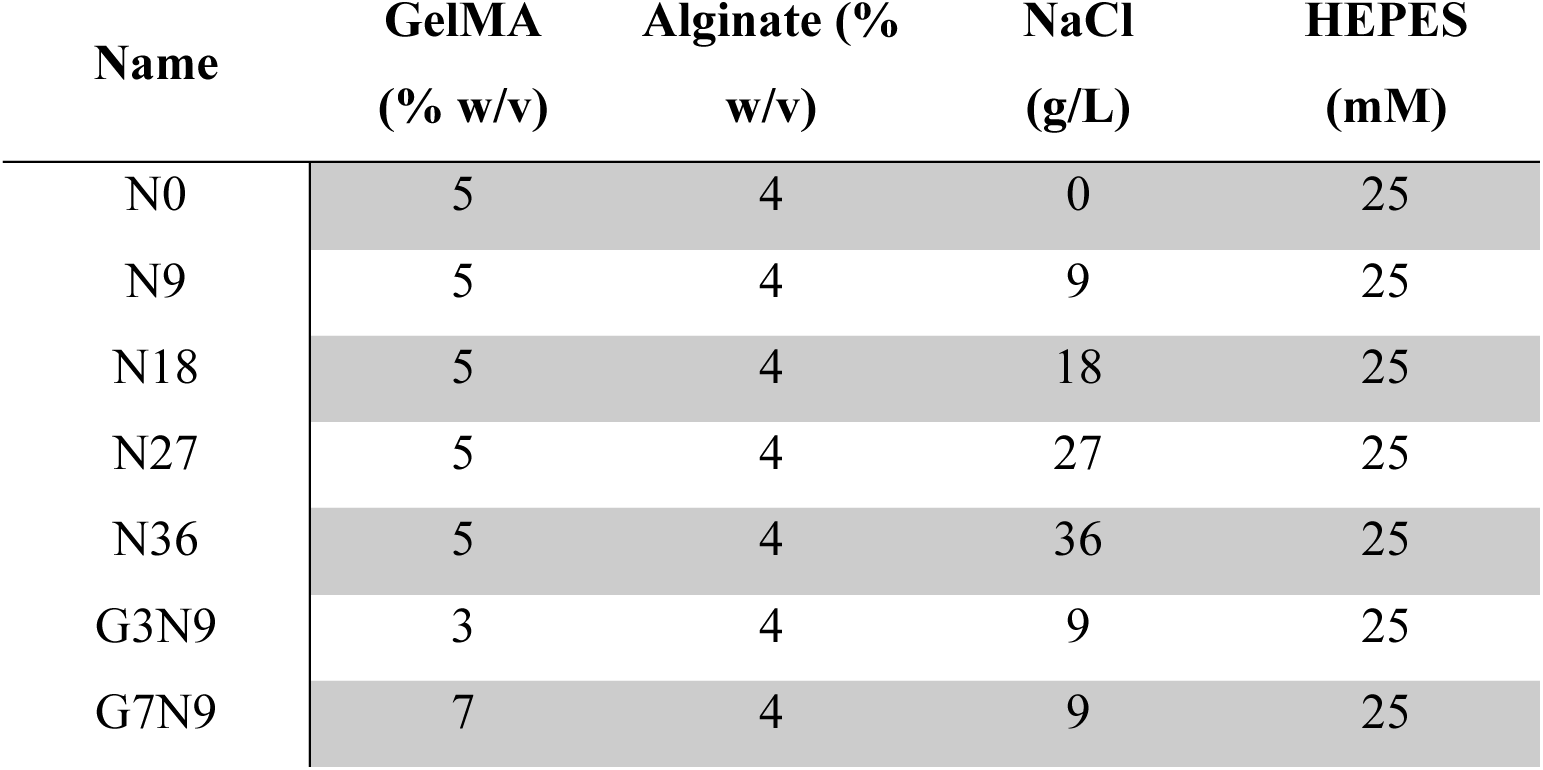
Formulation composition of ATPS inks.

### Image analysis and ATPS characterisation

The dispersed droplet phase at the various NaCl concentrations was observed inside the microfluidic chip with an inverted microscope (Zeiss - Observer.Z1) equipped with a high-speed camera (Photron FASTCAM Mini UX, 500 fps). A confocal microscope (Olympus IX83) was employed to image the porosities of the printed scaffolds at increasing NaCl concentrations. The autofluorescence of the alginate (405 nm)[55] enabled visualisation of the external and continuous phase of the samples. The percentage fraction of the area occupied by the internal phase was calculated using Equation (1):

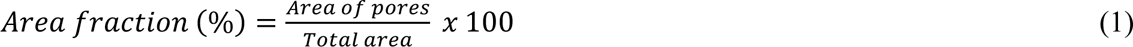

### ATPS scaffolds 3D printing

The ATPS phase was printed using a custom-made 3D bioprinter.[52] A MATLAB-generated g-code defined an 8mm x 8mm lattice scaffold (20 layers, 100 µm gap). Scaffold printing was performed by deposition on a 1mm-thick glass slide via a microfluidic printhead, allowing instantaneous cross-linking of the fibres by ionic interaction with CaCl_2_ (**Figure 1**). Lattice grids were 3D printed using ATPS inks at various concentrations of GelMA (3 – 7 %), alginate (4 %), Ru (1 mM) and SPS (10 mM) in NaCl (0 – 9 – 18 – 27 – 36 g/L) and HEPES (25 mM). Following deposition, the samples were cured (405 nm, 10 min), further cross-linked with a CaCl_2_ solution (0.33 M in NaCl (9 g/L) in HEPES (25 mM)) and stored in HBSS for analysis.

*Selective dissolution*: For the selective dissolution of GelMA, scaffolds were printed without Ru/SPS, cross-linked with CaCl_2_, and incubated in HBSS at 37 °C for 3 h to dissolve non-cross-linked GelMA. For the selective dissolution of alginate, following regular physical cross-linking of the alginate (CaCl_2_) and chemical curing of the GelMA (405 nm), the 3D printed samples were immersed overnight in a PBS-based solution with alginate lyase (0.33 mg ml^-1^). *Scanning Electron Microscopy*: Freeze-dried printed scaffold constructs (monophasic N0 and biphasic N36) were inspected by scanning electron microscopy (SEM) (HR-FESEM, AURIGA Zeiss model) working at 1.50 kV. The samples were first cut in half, then placed on aluminium stubs and metallised with a chromium plasma to enhance their conductivity. Images were taken at the following magnifications: 100x – 500x – 1000x.

### Optical coherence microscopy

To assess the difference in the architecture of the printed ATPS constructs, extreme conditions (N0 and N36) were evaluated by optical coherence tomography (OCT).[56] An OCT microscope (Thorlab OCT 4) with central wavelength of 880 nm and a sensitivity of 106 dB was operated with a 4x objective acquiring 3D images at 1024 x 1024 x 2048 pixels for the X, Y and Z axes with an imaging depth of 800 µm.

*Confocal imaging*: The internal and external structure of the printed ATPS were studied using a confocal laser scanning microscope. Prior to printing, GelMA-Rhodamine B and alginate-FITC were included in the different ATPS formulations. The scaffolds were then 3D printed, cross-linked, washed in HBSS and transferred into µ-Slide 8 Well (ibidi) for microscopic observation. A 559 nm laser was used for the observation of the phase of GelMA-Rhodamine B, and a 473 nm laser for the outer phase of alginate-FITC. The acquired images were processed with ImageJ software.

### Rheological measurements of emulsion inks

The rheological properties of the different ATPS formulations were analysed using a rotational rheometer (Anton Paar MCR 102) at 25 °C and 37 °C equipped with a cone-plate geometry (25 mm diameter, 1° cone angle). The shear viscosity behaviour of the ATPS solutions was evaluated by performing a shear rate from 1 to 9000 s^-1^.

### Dynamic mechanical testing of 3D printed ATPS scaffolds

The mechanical properties of the ATPS 3D constructs were measured using a DHR-2 controlled stress rotational Rheometer (TA Instruments, Waters) equipped with a parallel plate geometry (20mm diameter) and a Peltier plate system at 23 °C. The instrument was calibrated in DMA mode for uniaxial compression testing to determine storage (E’) and loss (E’’) moduli of hydrogel samples. Samples were pre-loaded at 0.4 N and frequency sweep tests (0.1 – 1 Hz) were performed at a constant displacement between 20 – 50 μm depending on the hydrogel linear viscoelastic region (LVR). Mean values and standard deviation were obtained by measuring three samples for each composition.

### In vitro degradation test

A degradation test was performed on scaffolds with both chemical curing of GelMA and ionic cross-linking of alginate, and on samples with only ionic cross-linking of alginate. The scaffolds were incubated in HBSS for 7 days at 37°C.[57] Prior to weighing, the samples were taken, dried and weighed at specific time points (day 0 - 1 - 3 - 7). *Swelling strands*: N0 and N36 scaffolds were printed and the variation in fibre dimension after 0 - 12 and 24 h was evaluated. The samples were printed and crosslinked both chemically (GelMA with Ru/SPS) and physically (alginate with CaCl_2_). The fibres were observed using an inverted microscope (AX10, Zeiss, Germany) with a 20x objective and the images were taken with an integrated camera (Axiocam 305 colour, Zeiss, Germany). The fibre diameters were measured at different time points with ImageJ software.

### Cell culture

The A549 human lung carcinoma epithelial-like cell line, C2C12 mouse pre-myoblast cell line and MG63 human osteosarcoma cell line were obtained from ATCC (USA). The cell lines were cultured in DMEM culture medium supplemented with 10% FBS, 100 U Penicillin, 0.1 mg ml^-1^ streptomycin and 2 mM L-glutamine. The cells were thawed, plated at 3×10^5^ cells/flask (75cm^2^, Corning®, NY, USA) and the medium was changed twice a week. Human bone marrow stromal cells (HBMSCs) were donated by Umberto I Hospital (Rome) and in accordance with the Declaration of Helsinki and its later amendments and approved on June 22, 2023 by the Institutional Review Board (Department of Molecular Medicine, Sapienza University of Rome, Italy). HBMSCs were cultured with αMEM with the same supplement as DMEM. The cells were thawed, plated at 5×10^5^ cells/flask (182 cm^2^, Corning®, NY, USA) and the medium was changed twice a week. All cells were incubated at 37 °C and 5% CO_2_.

### 3D Bioprinting

All biomaterials, solutions and printheads for ATPS-bioinks were sterilised via 30-min UV cycles for powders, filtration (0.22 µm syringe filter, ReliaPrep™, VWR) for solutions and flushing with EtOH first and PBS later inside the channels. Cells (A549, C2C12, MG63, HBMSCs) were detached (10 minutes, 37 °C, 5% CO_2_) using Trypsin-EDTA (1x) at 80-90 % confluence. The cells pellet was resuspended in culture medium and the biomaterial was added and mixed well prior to loading the bioink. The final cell concentration within the biomaterial was 5 million ml^-1^. Both the biomaterial, the cross-linking solution (CaCl_2_ 0.33 M in 80% glycerol) and washing solution (80% glycerol) were loaded into 3 sterile 3 ml syringes (Fisherbrand™, Fisher Scientific) and printing was performed in the absence of light to avoid premature cross-linking of the gelMA during scaffold fabrication. All syringes were set at a speed rate of 30 µl ml^-1^. The obtained scaffolds were exposed to visible light (10 min) and then to CaCl_2_ (10 min) for cross-linking. The grids containing A549, C2C12 and MG63 cell lines were cultured in DMEM, while those containing HBMSCs were cultured in αMEM. For osteogenesis studies, αMEM was supplemented with osteogenic differentiation factors: AA2P (100 µM), β-GP (10 mM), dexamethasone (10 nM).

### Cell viability

Live/Dead assay was performed by adding 0.4 ug ml^-1^ of Calcein and 0.2 ug ml^-1^ of Propidium Iodide to FBS-free medium. The solution was then added on top of the previously washed (HBSS) scaffolds until complete coverage and incubated in the dark for 30 minutes (37 °C; 5% CO_2_). Subsequently, the scaffolds were washed in HBSS and analysed with a confocal microscope (Olimpus IX83). Cell viability was quantitatively evaluated by ImageJ software analysis.

### Immunofluorescence

The scaffolds were fixed with 4% PFA, permeabilised with 0.1% Triton X-100 and then immersed in 1% BSA to block autofluorescence of the biomaterials. Samples were incubated overnight at 4 °C in the dark with primary antibodies (anti-Rabbit, 4 µg ml^-1^; anti-Mouse, 2 µg ml^-1^) in 1% BSA, followed by secondary antibodies staining. For BMP-2 staining: BMP-2 antibody (5 µg ml^-1^) and DAPI (1:1000). For phalloidin staining, samples were incubated with AlexaFluor™ 488 phalloidin (1:40) and DAPI (1:1000) after the autofluorescence blocking step. The scaffolds were observed with a 40x objective under a laser scanning confocal microscope.

### Alkaline Phosphatase staining

Scaffolds were washed twice in HBSS, followed by 95% EtOH and a final HBSS rinse. After drying, samples were incubated in a HBSS-based solution containing Naphthol AS-MX Phosphate Alkaline Solution and Fast Red Violet LB Salt for 60 min at 37 °C in the dark. Following three HBSS washes, scaffolds were imaged using an inverted microscope with a 20x objective with an integrated camera.

### Alizarin Red S staining

ATPS constructs were first removed from the culture medium, washed with HBSS and then fixed for 1 h in 4% PFA in HBSS. Subsequently, the samples were embedded in OCT at - 80 °C, then cryosectioned at −20 °C (8 µm thickness, Leica CM1850, Leica Biosystems) and mounted on SuperFrost glass slides (Epredia™ SuperFrost Plus™). The slices were stained with a previously prepared Alizarin Red S solution. Briefly, 3.5 g of Alizarin Red S were dissolved in 150 ml DW. The pH was adjusted to 4.1-4.3 using 0.5% ammonium hydroxide and the volume was finally brought to 250 ml by monitoring the pH. The slices were soaked in Alizarin Red S solution for 1 hour, then washed with DW and visualised with an inverted microscope.

### Quantitative reverse transcription polymerase chain reaction

The expression of specific genes related to osteogenic differentiation (**Table 4**) of HBMSCs encapsulated in ATPS hydrogels was determined by quantitative reverse transcription polymerase chain reaction (RT-qPCR). The samples were first degraded in a solution containing 1 mg ml^-1^ collagenase D and 0.33 mg ml^-1^ alginate lyase. RNA was isolated via TRIzol Reagent, extracted (RNeasy Mini Kit, QIAGEN) and quantified using a nanodrop spectrophotometer (NanoDrop 2000c, ThermoFisher Scientific). RNA was converted to cDNA using the iScript™ cDNA Synthesis Kit (Bio-rad). RT-qPCR was performed using iTaq™ Universal SYBR® Green Supermix (Bio-rad) and gene expression quantified using a Real-Time PCR System (Applied Biosystems ViiA 7, ThermoFisher Scientific). The cell pellet collected at day 0 was used as a control. The expression of targeted genes related to osteogenic differentiation was calculated by threshold cycle (ΔΔCt) method, normalised to GAPDH as housekeeping reference gene and reported as fold change (2^-ΔΔCt^).

**Table 4.**
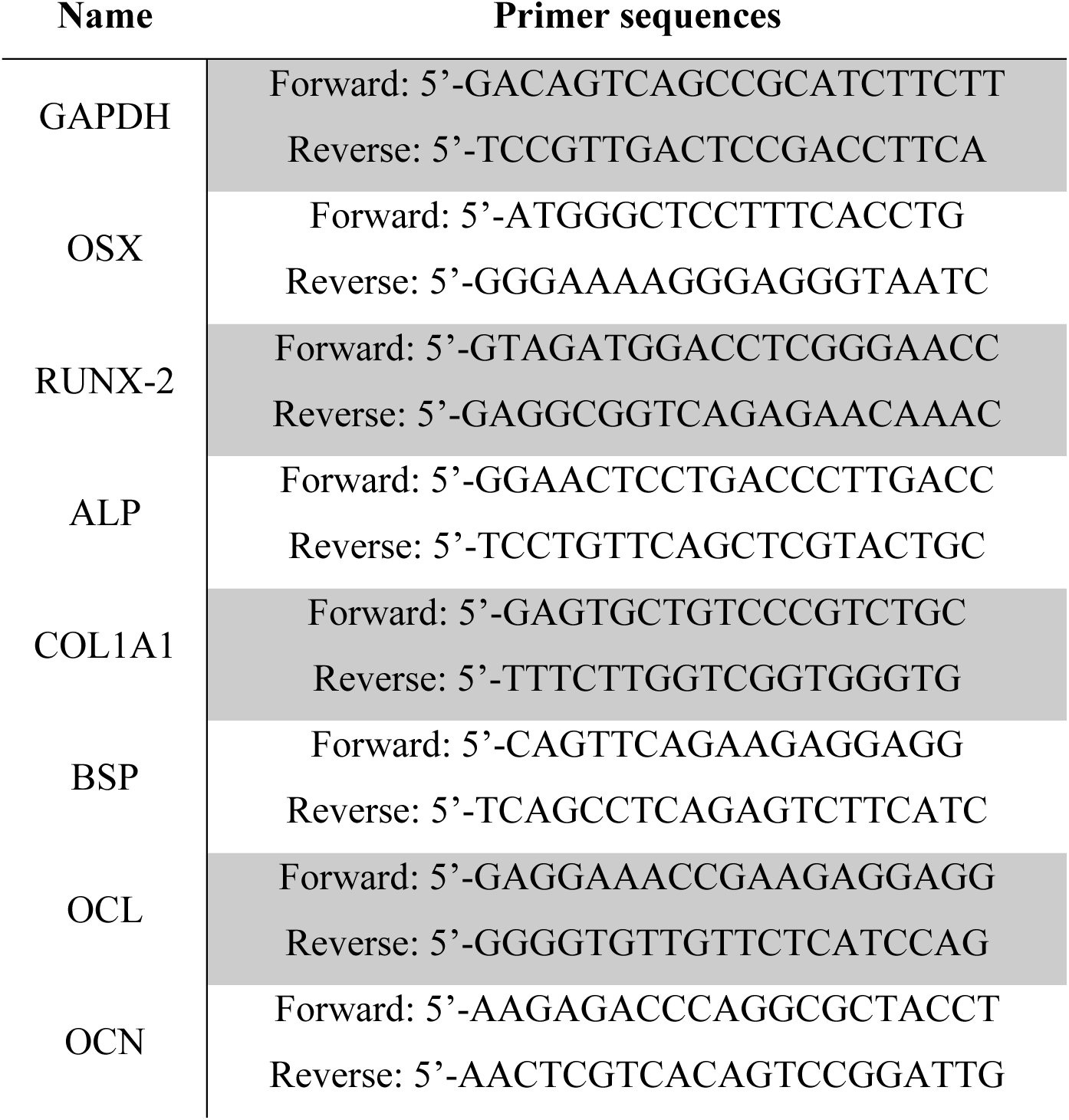
Primer sequences used for RT-qPCR.

### Chick chorioallantoic membrane assay

All animal procedures were performed in accordance with European Directive 2010/63/EU and Italian Legislative Decree No. 26 of 4 March 2014. Following a previously reported protocol,[58] on day 0 fertilised chicken eggs were placed in a Cimuka CT120SH incubator (Ankara, Turkey) and maintained under controlled conditions at 37.7°C and 50% relative humidity. Eggs were rotated 90 degrees every 3.2 hours during the initial incubation period. On day 7, a 1 cm² opening was made in the eggshell using a sterile scalpel. In addition, scaffolds of identical composition were pre-soaked in a solution containing 10 μg VEGF in 1 mL HBSS prior to implantation. Each scaffold was carefully inserted through the eggshell window in a planar configuration and precisely positioned on the chick chorioallantoic membrane (CAM). The eggs were then sealed with sterile parafilm tape, incubated in a stationary position and monitored daily. On day 14, the scaffolds were removed and their integration with the CAM was assessed using a stereomicroscope equipped with a digital camera. Finally, the experimental procedure was terminated in accordance with established ethical guidelines.

### AI-based analysis of vascular networks

An image-based analysis was used to evaluate the vascular architecture of the CAM in the presence or absence of 3D bioprinted constructs. Images (n=3 per condition) were captured using a Zeiss Stemi 305 stereomicroscope at 1x magnification. The dataset included three groups of experimental conditions: -VEGF (N36 and N0 inks were combined with HBMSCs), +VEGF (same inks formulation with HBMSCs and a filler material supplemented with 10 µg/mL VEGF) and empty (CAM without implanted constructs). The analytical workflow consisted in the assessment of angiogenic response using the Chalkley Score (CS). The CS was automatically calculated in the user interface. The user has to draw a box to locate the biographical construct. The number of circles touching the vessels in the image was then counted. The CS was calculated by rotating the grid from 0° to 360°, with an increment of 10° for each rotation. Only the ten highest CS values were considered for each image.

### Statistical analysis

Statistical analysis was performed via GraphPad Prism (Graph Pad Software Inc., La Jolla, CA). The D’Agostino-Pearson normality test was applied to assess variations in the data. One-way ANOVA and two-way ANOVA were used to evaluate differences between groups. The sample size (n) is shown in the figure captions. Significantly different data were selected as p < 0.05.

## Acknowledgements

G.C. acknowledges funding from MTF Biologics (OSTEOMIMIC). This research was partially funded by grants from ERC-2019-Synergy Grant (ASTRA, n. 855923); EIC-2022-PathfinderOpen (ivBM-4PAP, n. 101098989); Project “National Center for Gene Therapy and Drugs based on RNA Technology” (CN00000041) financed by NextGeneration EU PNRR MUR—M4C2—Action 1.4—Call “Potenziamento strutture di ricerca e creazione di “campioni nazionali di R&S” (CUP J33C22001130001). A.B. and R.A. acknowledge MIUR for financial support (Project PRIN 2022ZA77J2 ICARUS – CUP B53D23009010006). Cartoons in figures were created with BioRender.

## Table of contents

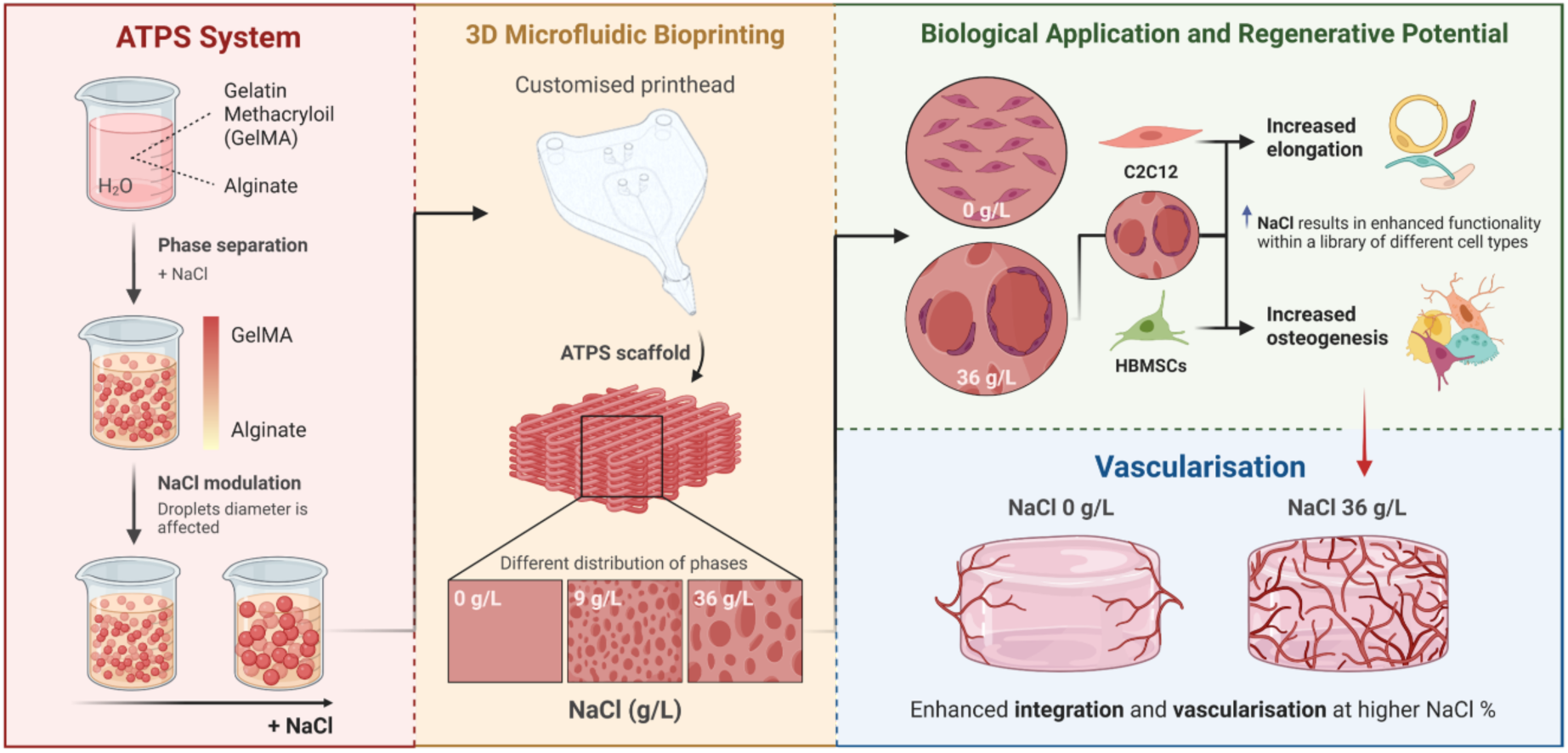

Aqueous two-phase systems (ATPS) enable the formation of biomimetic interfaces crucial for tissue engineering. However, clinical translation remains limited by the challenge of precisely controlling cellular compartmentalisation. Here, we developed ATPS biomaterial inks for 3D bioprinting allowing tuneable droplet formation via NaCl modulation. This strategy promoted controlled cell patterning, osteogenic differentiation and vascular integration, offering a promising approach for regenerative medicine applications.

## Supporting Information

**Figure S1.**
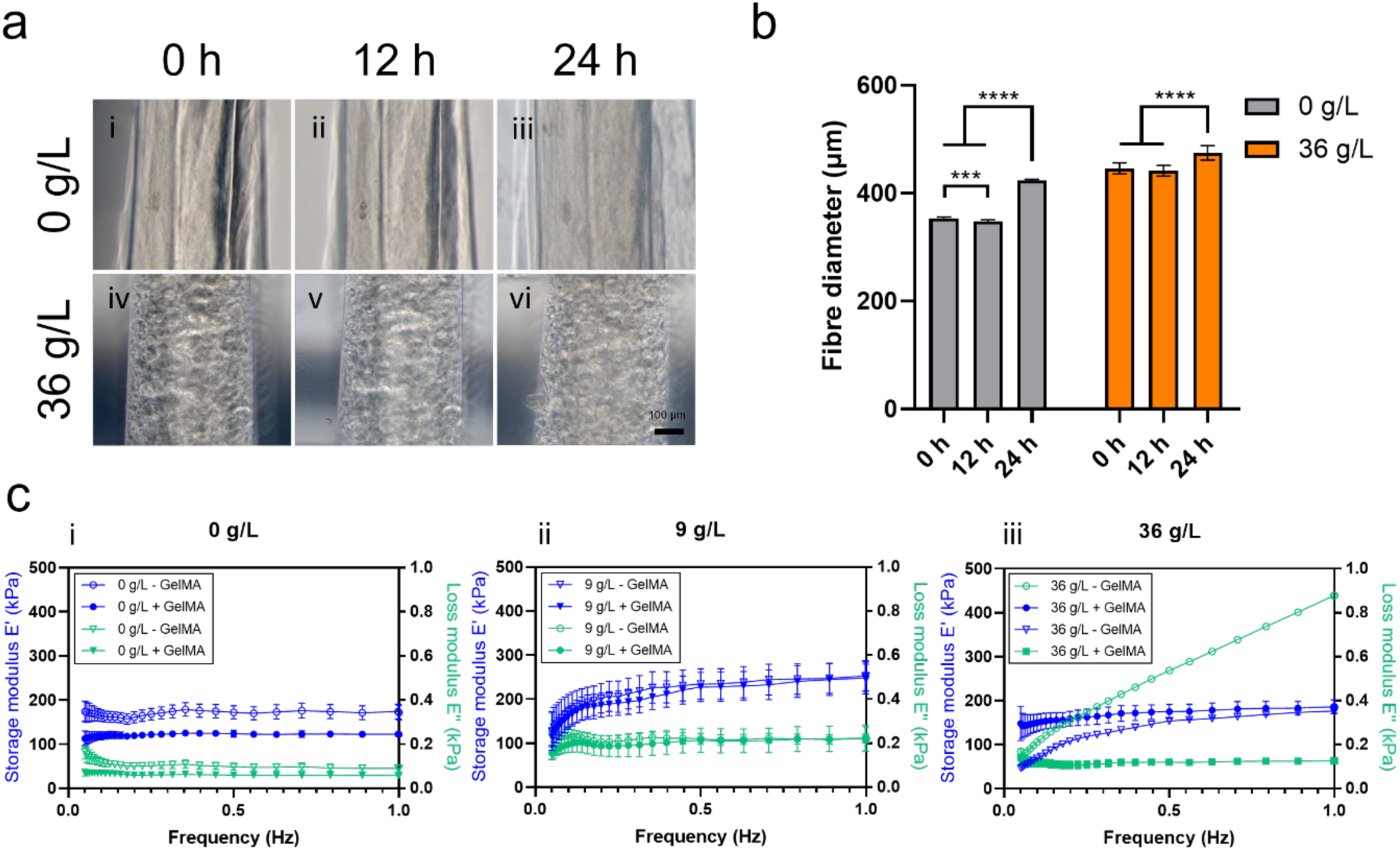
Swelling and mechanical analysis of printed ATPS fibres. a) Images of fibres swelling at 0 g/L NaCl taken a-i) 0 hours, a-ii) 12 hours, a-iii) 24 hours and fibres at 36 g/L NaCl taken a-iv) 0 hours, a-v) 12 hours and a-vi) 24 hours. b) Quantification of printed fibres swelling at 0 - 36 g/L NaCl. c) E’ and E’’ values obtained from compression tests performed on samples at c-i) 0, c-ii) 9 and c-iii) 36 g/L with and without GelMA cross-linking. Scale bar: (a) 100 μm. Statistical significances were assessed by one-way ANOVA. Mean ± S.D. n=3, ****p<0.0001, ***p<0.001.

**Figure S2.**
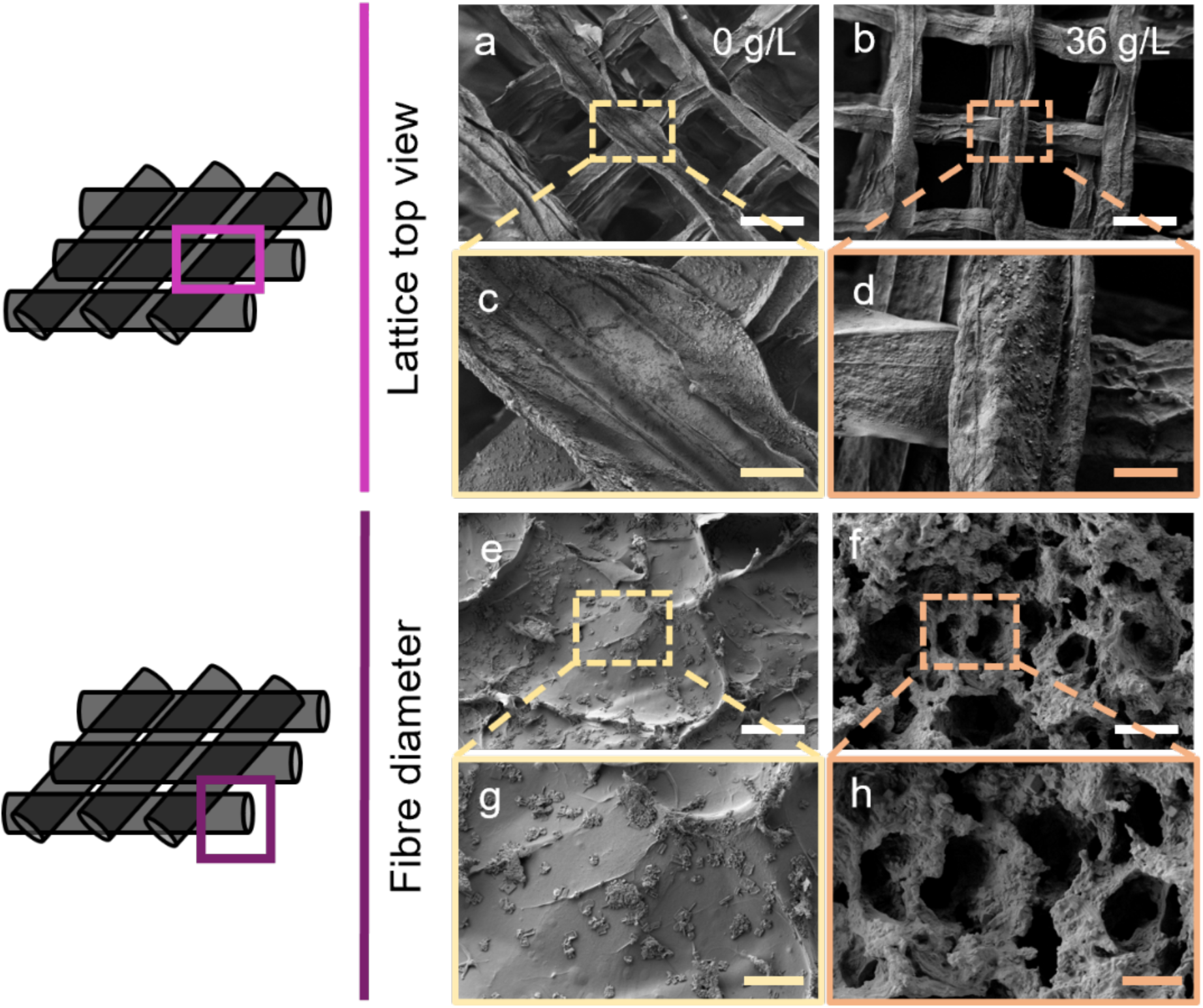
SEM characterisation of ATPS scaffolds. SEM top view images of the freeze-dried fibres at i, ii) 0 g/L NaCl and iii-iv) 36 g/L NaCl. SEM images of sections of lyophilised fibres at v-vi) 0 g/L NaCl and vii-viii) 36 g/L NaCl. Scale bars: (a,b) 500 μm, (c,d) 100 μm, (e,f) 100 μm, (g,h) 50 μm. Mean ± S.D. n=3.

**Figure S3.**
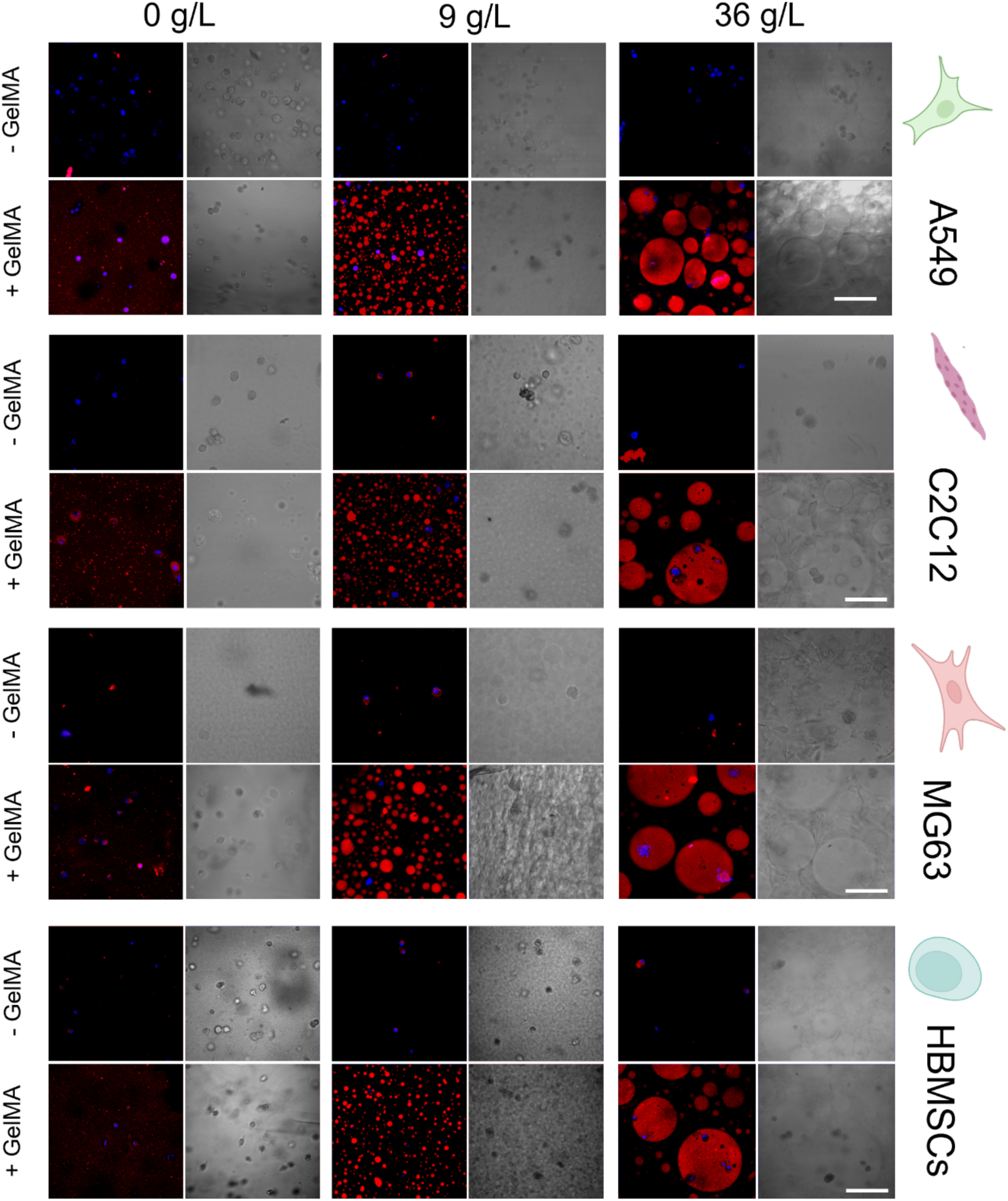
Partitioning of different cell types within the various ATPS scaffold formulations. Brightfield and confocal (merged) images representative of cells labelled with DAPI to show the nucleus (blue) of the different cell types (A549, C2C12, MG63, HBMSCs), and GelMA marked with rhodamine B to show the inner phase (red) of the scaffolds at 0 - 9 - 36 g/L. In all cell types, images were taken both under conditions where GelMA was not chemically cross-linked (-GelMA) and under conditions where GelMA was chemically cross-linked (+GelMA). Scale bars: (a, b, c, d) 100 μm. Mean ± S.D. n=3.

**Figure S4.**
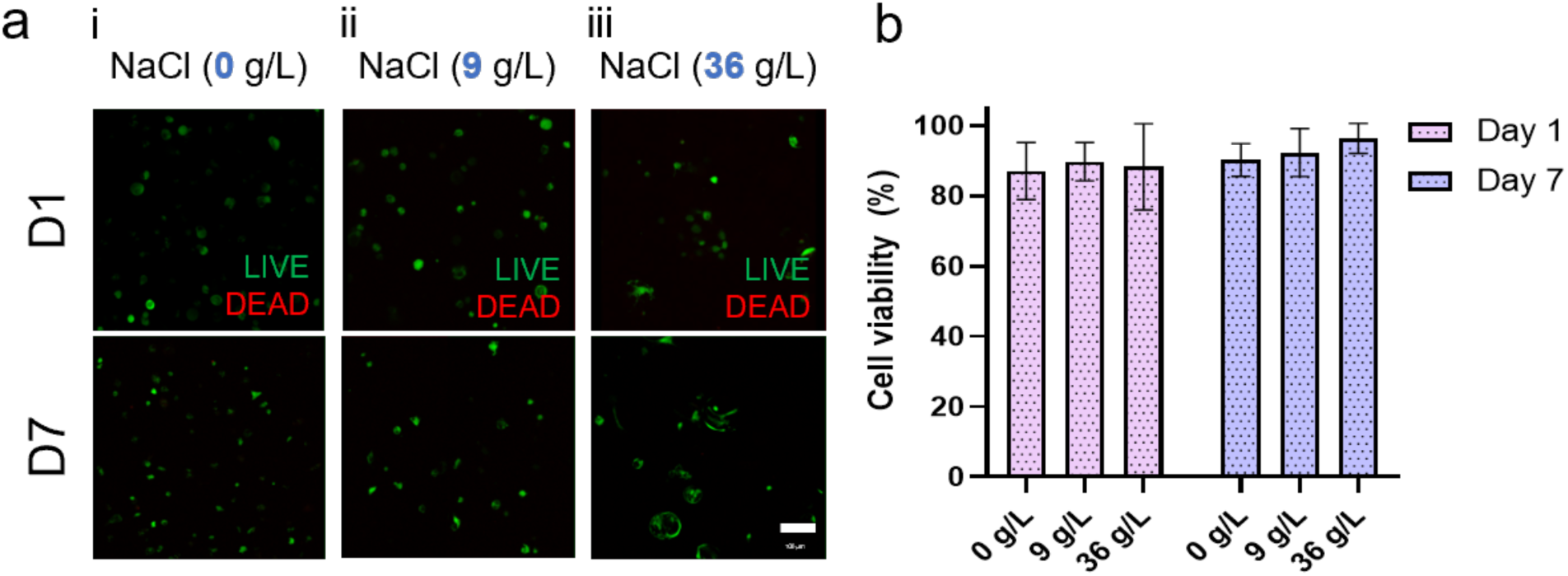
Cell viability of HBMSCs encapsulated in ATPS scaffolds. a) Confocal images with HBMSCs cells labelled with Calcein to show live cells (green) and cells labelled with propidium iodide to show dead cells (red) for the conditions a-i) 0 g/L NaCl, a-ii) 9 g/L NaCl and a-iii) 36 g/L NaCl. b) Quantification of cell viability at day 1 and day 7 for samples 0 - 9 - 36 g/L. Scale bar: (a) 100 μm. Statistical significances were assessed by two-way ANOVA. Mean ± S.D. n=3.

