## Supplementary Info for "Aqueous two-phase bioinks for discrete packing and compartmentalisation of 3D bioprinted cells"

### Supporting Information

**Aqueous two-phase bioinks for discrete packing and compartmentalisation of 3D bioprinted cells**

*Martina Marcotulli, Arianna Iacomino, Federico Serpe, Lucia Iafrate, Marco Bastioli, Giorgia Montalbano, Biagio Palmisano, Silvia Franco, Roberta Angelini, Alessandro Corsi, Mara Riminucci, Giancarlo Ruocco, Chiara Scognamiglio, Andrea Barbetta*, and Gianluca Cidonio**

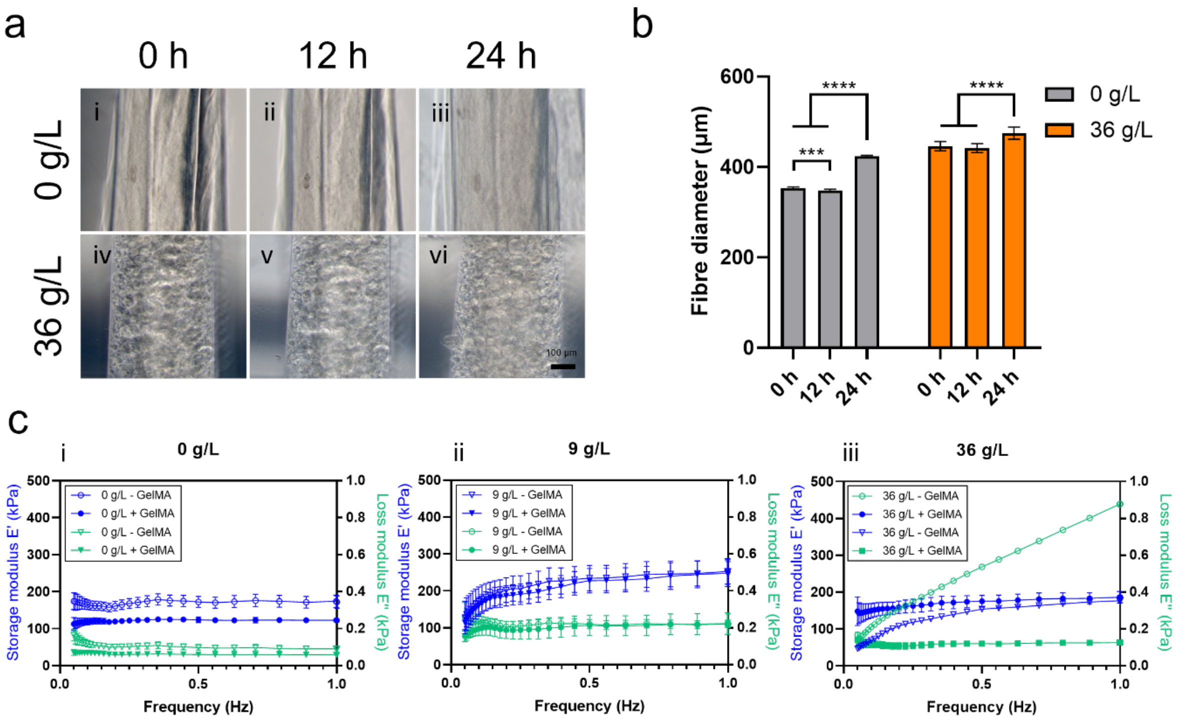


**Figure S1**. Swelling and mechanical analysis of printed ATPS fibres. a) Images of fibres swelling at 0 g/L NaCl taken a-i) 0 hours, a-ii) 12 hours, a-iii) 24 hours and fibres at 36 g/L NaCl taken a-iv) 0 hours, a-v) 12 hours and a-vi) 24 hours. b) Quantification of printed fibres swelling at 0 - 36 g/L NaCl. c) E' and E" values obtained from compression tests performed on samples at c-i) 0, c-ii) 9 and c-iii) 36 g/L with and without GelMA cross-linking. Scale bar: (a) 100 µm. Statistical significances were assessed by one-way ANOVA. Mean ± S.D. n=3, ****p<0.0001, ***p<0.001.

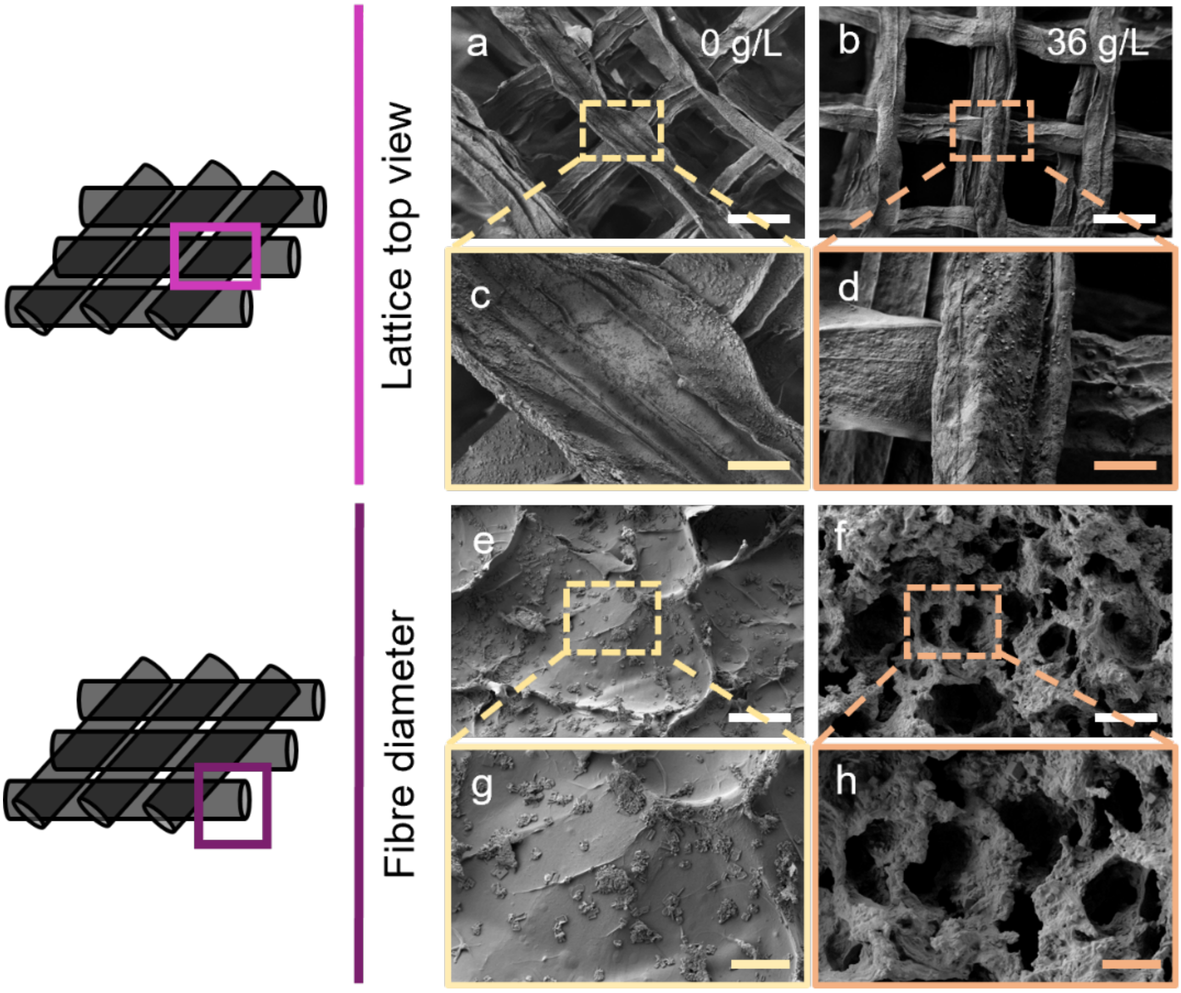


**Figure S2**. SEM characterisation of ATPS scaffolds. SEM top view images of the freeze-dried fibres at i, ii) 0 g/L NaCl and iii-iv) 36 g/L NaCl. SEM images of sections of lyophilised fibres at v-vi) 0 g/L NaCl and vii-viii) 36 g/L NaCl. Scale bars: (a,b) 500 µm, (c,d) 100 µm, (e,f) 100 µm, (g,h) 50 µm. Mean ± S.D. n=3.

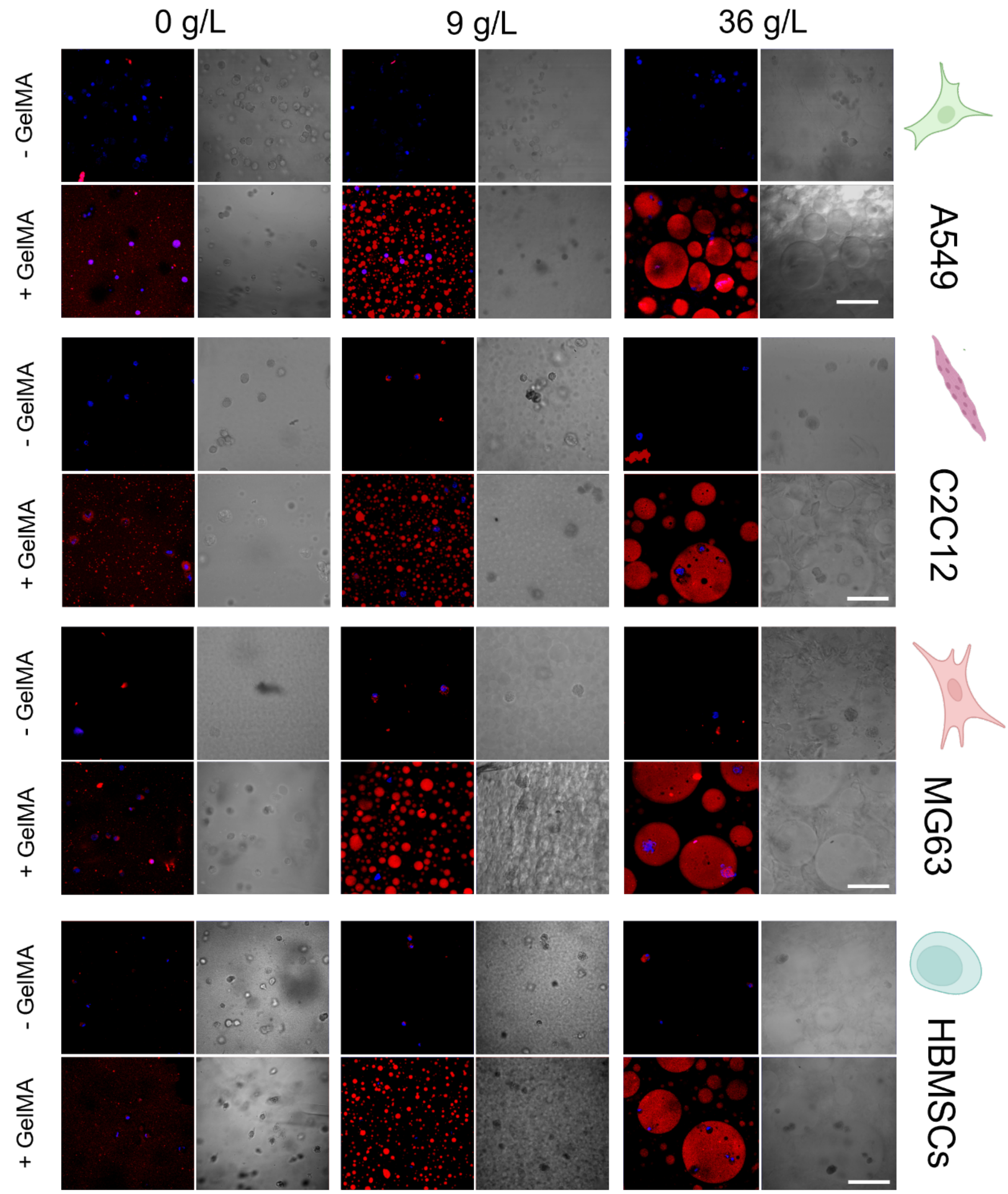


**Figure S3**. Partitioning of different cell types within the various ATPS scaffold formulations. Brightfield and confocal (merged) images representative of cells labelled with DAPI to show the nucleus (blue) of the different cell types (A549, C2C12, MG63, HBMSCs), and GelMA marked with rhodamine B to show the inner phase (red) of the scaffolds at 0 - 9 - 36 g/L. In all cell types, images were taken both under conditions where GelMA was not chemically cross-linked (-GelMA) and under conditions where GelMA was chemically cross-linked (+GelMA). Scale bars: (a, b, c, d) 100 µm. Mean ± S.D. n=3.

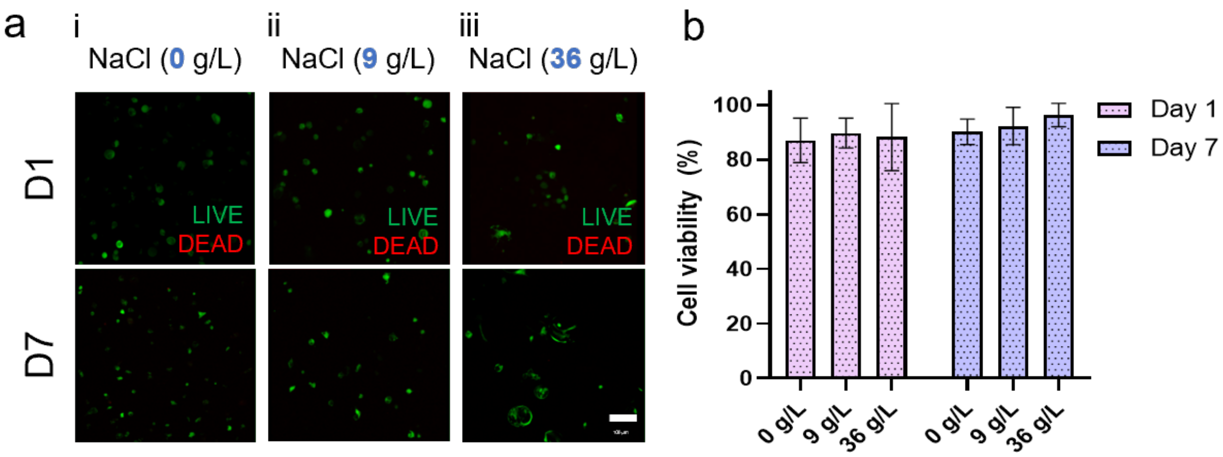


**Figure S4**. Cell viability of HBMSCs encapsulated in ATPS scaffolds. a) Confocal images with HBMSCs cells labelled with Calcein to show live cells (green) and cells labelled with propidium iodide to show dead cells (red) for the conditions a-i) 0 g/L NaCl, a-ii) 9 g/L NaCl and a-iii) 36 g/L NaCl. b) Quantification of cell viability at day 1 and day 7 for samples 0 - 9 - 36 g/L. Scale bar: (a) 100 µm. Statistical significances were assessed by two-way ANOVA. Mean ± S.D. n=3.
